# Detecting oncogenic selection through biased allele retention in The Cancer Genome Atlas

**DOI:** 10.1101/2020.07.03.186593

**Authors:** Juliet Luft, Robert S. Young, Alison M. Meynert, Martin S. Taylor

**Affiliations:** MRC Human Genetics Unit, MRC Institute for Genetics and Molecular Medicine, University of Edinburgh, Crewe Road, Edinburgh, EH4 2XU, UK; Centre for Global Health Research, Usher Institute, University of Edinburgh, Teviot Place, Edinburgh, EH8 9AG, UK

## Abstract

**Background:** The loss of genetic diversity in segments over a genome (loss-of-heterozygosity, LOH) is a common occurrence in many types of cancer. By analysing patterns of preferential allelic retention during LOH in approximately 10,000 cancer samples from The Cancer Genome Atlas (TCGA), we sought to systematically identify genetic polymorphisms currently segregating in the human population that are preferentially selected for, or against during cancer development.

**Results:** Experimental batch effects and cross-sample contamination were found to be substantial confounders in this widely used and well studied dataset. To mitigate these we developed a generally applicable classifier (GenomeArtiFinder) to quantify contamination and other abnormalities. We provide these results as a resource to aid further analysis of TCGA whole exome sequencing data. In total, 1,678 pairs of samples (14.7%) were found to be contaminated or affected by systematic experimental error. After filtering, our analysis of LOH revealed an overall trend for biased retention of cancer-associated risk alleles previously identified by genome wide association studies. Analysis of predicted damaging germline variants identified highly significant oncogenic selection for recessive tumour suppressor alleles. These are enriched for biological pathways involved in genome maintenance and stability.

**Conclusions:** Our results identified predicted damaging germline variants in genes responsible for the repair of DNA strand breaks and homologous repair as the most common targets of allele biased LOH. This suggests a ratchet-like process where heterozygous germline mutations in these genes reduce the efficacy of DNA double-strand break repair, increasing the likelihood of a second hit at the locus removing the wild-type allele and triggering an oncogenic mutator phenotype.

## Introduction

Loss-of-heterozygosity (LOH) describes the somatic loss of genetic material from one copy of a heterozygous locus. It can occur as a consequence of whole or partial chromosome deletion, or as a copy-neutral event, in which one copy is replaced by the other - for example through homologous repair^1^ or locus duplication followed by loss of the non-duplicated allele. LOH often occurs as the ‘second-hit’ in tumour initiation, where somatic loss of the wild-type (WT) copy opposite either a germline or somatic mutation drives cancer progression^2,3^.

Previous studies of LOH have sought to identify novel tumour suppressor genes by mapping patterns of LOH in tumours, but were hampered by low-resolution data and inadequate sample sizes^4^. A more recent study in ovarian cancer overlapped recurrent regions of LOH with somatic mutation data, and whilst their results revealed strong selection of known cancer genes (deletion of the WT allele in 94% of cases with deleterious somatic TP53 or BRCA1 mutations), it failed to reveal novel drivers^5^. In contrast, studies in cutaneous squamous cell carcinoma, ovarian cancer and colorectal cancer revealed evidence of preferential allelic imbalance of putative germline risk variants^5–7^, indicating that LOH may also have a role in the selection of small-effect, inherited, cancer-predisposing variants. By systematically quantifying biased allele loss or retention across a large cohort of whole exome sequencing (WXS) data, we sought to explore genetic selection of common cancer-associated variants during cancer progression.

The Cancer Genome Atlas (TCGA) is a public resource of genomic, clinical and associated data from over 10,000 patients across 36 types of cancer, including WXS from matched tumour:normal sample pairs^8^. This wealth of data is extensively used and a valuable resource in the field of cancer genomics, but is subject to the influence of batch effects and technical artefacts^9–12^. To control for these effects we performed a systematic analysis of mapping and sequencing artefacts, exome target capture kit biases and contamination in TCGA, and developed a workflow to identify and remove these confounding influences. Strikingly we found evidence of contamination and other issues in 1,678 pairs of samples (14.7%), a result that may have had unforeseen impact on previous analyses performed using this dataset.

After rigorous filtering of the data, we did not identify preferential retention of common variants via LOH, although we did observe an overall trend for biased retention of cancer-associated risk alleles identified by genome wide association study (GWAS). Subsequent targeted analysis revealed strong oncogenic selection of predicted damaging germline variants in recessive tumour suppressor genes and preferential retention of predicted damaging germline variants in 25 protein interaction pathways, predominantly those involved in DNA damage repair and cell cycle. Of 1,175 patients with a predicted damaging germline variant in a recessive tumour suppressor gene, 284 had LOH with retention of the damaging allele at the locus (24.17%; 2.96% of 9,602 total patients analysed).

## Results

### Initial analysis of biased allele retention during loss-of-heterozygosity in tumours

Analysis of biased LOH was performed using paired normal and primary tumour samples from 9,905 patients across 36 cancer subtypes. Germline variants were called using Strelka2^13^ and GATK HaplotypeCaller^14^, and subsequent LOH calling was performed using CloneCNA^15^. To quantify allele retention bias during loss-of-heterozygosity, Fisher’s Exact tests were performed comparing reference versus alternative allele bias in samples that had undergone LOH versus those that hadn’t (Materials and Methods). Of the 210,456 loci tested (heterozygous in >= 50 normal samples), 74 variants had evidence of significantly biased retention of either the reference or alternative allele (Figure 1a; Fisher’s-exact test; Bonferroni corrected p < 0.05).

**Figure 1:**
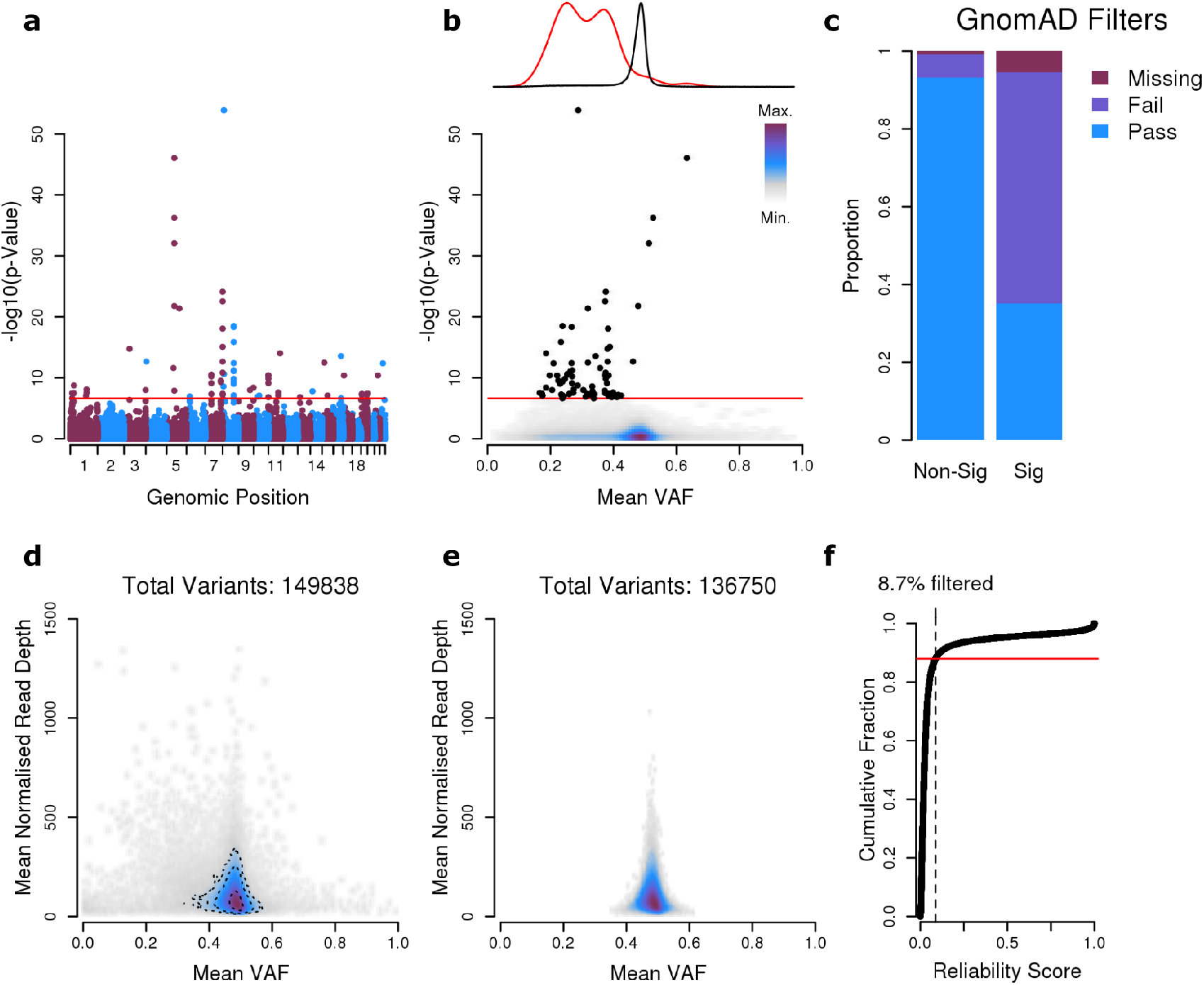
Artefactual germline variants influence loss-of-heterozygosity bias analysis. **a,** Pan-cancer LOH bias analysis. The Y-axis shows the −log10(p-value) from a Fisher’s exact test for LOH bias. The red line indicates the threshold for significance (Bonferroni corrected p < 0.05). **b,** Mean VAF in the normal sample of germline heterozygous individuals for each variant included in the LOH bias analysis, versus significance in the LOH bias analysis, colour indicates density of points. Individual points above significance threshold plotted in black. Curves above the plot show VAF density of non-significant variants (black) and significant variants (red). **c,** Proportion of variants identified as non-significant (Non-Sig) / significant (Sig) in the LOH bias analysis that pass / fail gnomAD variant filters, or are missing from the gnomAD database. **de**, Distribution of mean VAF versus mean normalised read depth from normal samples in (**d**) all variants and (**e**) variants that pass our filter. Colour indicates density of points. Contour lines in **d** show 50%, 75% and 90% limits. Only variants that appear heterozygous in at least 1% of TCGA normal samples are included. Variants that fail gnomAD filters or are missing from the database have been removed. **f,** Cumulative distribution plot showing the distribution of ‘Reliability’ scores across all variants. Red horizontal line indicates the filtering threshold used in this analysis (0.88).

Autosomal germline variants are expected to have heterozygous variant allele frequency (VAF; proportion of reads mapping to the non-reference allele) of approximately 50% in the normal samples. Investigation of the significantly LOH-biased variants revealed them to have a significantly lower VAF in the normal sample than non-significant common variants (heterozygous in >=1% of normal samples) (Figure 1b; mean VAF 1.5 fold lower, t-test, p = 2.5e-23). After cross-referencing our data with gnomAD^16^, a database of human genetic variation that includes the normal samples within TCGA; we found that the proportion of significantly LOH-biased variants failing gnomAD variant filters or missing from the gnomAD database was higher than seen for non-significant variants (Figure 1c; OR = 25.33, Fisher’s exact test, p-value = 1.1e-37). These results indicate that many of the significant allele retention biases we detected were the result of artefactual variants. Motivated by these results we undertook a systematic interrogation of TCGA WXS data covering three broad areas: 1) mapping and sequencing artefacts, 2) exome target capture kit biases and 3) sample specific abnormalities including contamination.

### Systematic detection of artefactual and unreliable germline heterozygous variants

We first removed all variants that failed the gnomAD filters or were missing from the gnomAD database, therefore focusing our analysis on consensus, high-quality germline variants. We found that the genomic regions with consistently low VAF giving rise to likely false biased allele retention signals, typically fall into one of two categories. First, lower alternative allele read-mapping rates in haplotype segments with clusters of non-reference alleles (example locus shown in Supplementary Figure 1a, associated normal sample VAF data in Supplementary Figure 1b,c). And second, in regions that have high sequence-identity paralogous regions elsewhere in the genome (example locus shown in Supplementary Figure 1d, associated tumour:normal VAF data in Supplementary Figure 1e).

Low VAF can often occur due to stochastic sampling of reads at low coverage and is not always indicative of an underlying problem; as such, a standard VAF thresholding measure is not enough to identify error-prone or hard-to-map loci. Consequently we developed a binomial-test based reliability score to predict the likelihood of a variant being ‘true’ given its observed VAF and read-depth across all normal heterozygous samples (Materials and Methods, Figure 1d). In total, 13,088 variants (8.7% of common variants) failed our threshold, and were subsequently removed (Figure 1e,f) - this included all 26 variants previously identified as significant in the LOH bias analysis, which passed gnomAD filters.

### Influence of exome target capture kit on the output of whole exome sequencing data

At many loci, samples sequenced by the same exome target capture kit tended to cluster by VAF and read depth (Figure 2a), demonstrating a kit specific effect on the raw sequence data, consistent with prior observations^17,18^. Additionally, we found that on average, individuals sequenced by the same kit had significantly more variants in common than those sequenced using a different kit (mean correlation coefficient 1.2 fold higher, t-test, p-value < 2.2e-16; Figure 2b,c; Supplementary Figure 2).

**Figure 2:**
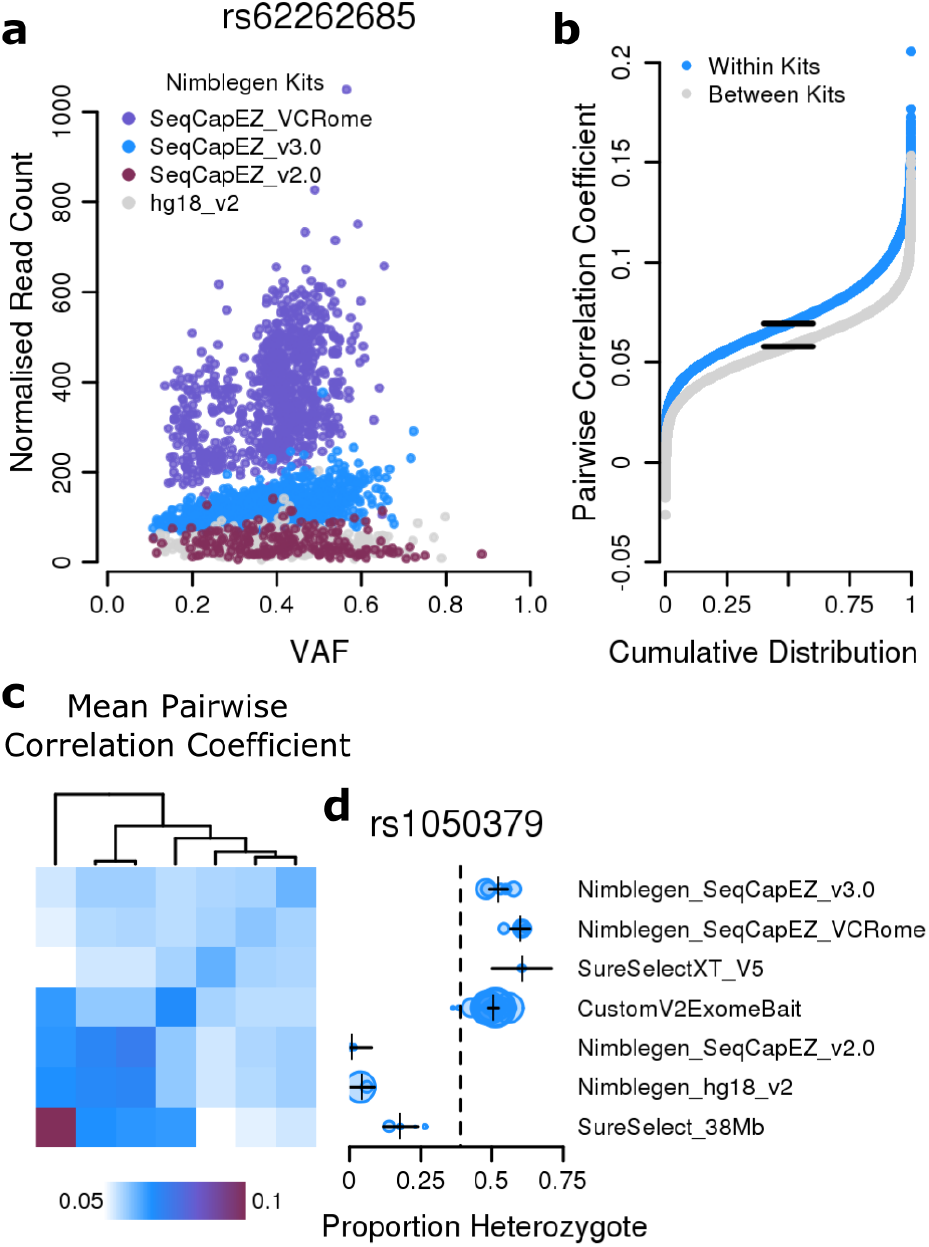
Exome target capture kit biases influence raw variant data and result in batch specific artefactual variant calls. **a,** Distribution of VAF versus normalised read depth at an example variant (rs62262685) for heterozygous normal samples, sequenced by the indicated exome target capture kit. To highlight the extent of variation within a single manufacturer, only kits from Nimblegen have been included though similar results are observed for other manufacturers. **b,** Correlation coefficients of pairs of patients sequenced by the same kit or by different kits. See extended methods for details, full results for all kit pairs in Supplementary Figure 1. **c,** Heatmap shows mean correlation coefficient between patients from each pair of kits, rows and columns are ordered as in **d**. **d,** Illustrative example of the filtering methodology. Plot shows proportion of white/European individuals heterozygous for an example variant (rs1050379), grouped by exome target capture kit. Points show individual cancer subtypes. Point sizes are proportional to the number of patients within each group. Vertical dashed line indicates the expected heterozygous frequency (hetFreq), calculated from the gnomAD non-Finnish European genome allele frequency (NFE_AF). Black crosses show the estimated effect of the exome target capture kit on the hetFreq, as calculated by linear regression; horizontal error bars = 2*SE. All kits significant p < 0.001.

We developed a linear regression based method to identify variants that were over-enriched in calls from specific kits (Figure 2d). This compared observed heterozygote frequency with a gnomAD derived allele frequency and stratified by both exome target capture kit and cancer subtype (Materials and Methods). In total, 80 variants (0.06%) were identified that passed previous filters and were significantly over-represented in one or more kits (Supplementary Figure 3c; Bonferroni corrected p < 0.05). Many previously filtered variants were significantly over-represented across all kits, although a small proportion showed a variable pattern of enrichment across the kits (Supplementary Figure 3a,b), indicating that some of the previously observed sequencing artefacts may be kit specific.

### Evidence of contamination in samples from The Cancer Genome Atlas

The final step of our interrogation of technical artefacts in TCGA WXS data was to investigate sample specific abnormalities. The VAF distribution of germline heterozygous variants in matched tumour:normal samples can be used to infer characteristics of the tumour - such as the fraction of the genome subject to LOH (Figure 3a,b), and the cellularity of the sample (Figure 3c). By quantifying deviations from the expected distribution, we can identify other abnormalities - such as contamination with genetic material from a different individual (Figure 3d-g) and low quality sequencing data (Figure 3h,i). Using a combination of metrics derived from the tumour:normal VAF distribution we developed a two step pipeline to identify and filter problematic samples (Extended Methods). First, hard thresholds were used to exclude samples with severely abnormal distributions (subsequently denoted: *X*); secondly, an ordinal logistic regression based classifier was trained to quantify the extent of contamination in each tumour:normal sample pair, and assign them to one of four qualitative groups: *C0* = non-contaminated, *C1* = minor contamination, *C2* = moderate contamination, *C3* = severe contamination (Figure 4a-d). Applying our pipeline to the TCGA samples, 185 (1.6%) pairs of samples were excluded in the first step, and 1,493 (13.3%) pairs of the remaining samples were identified as contaminated (*C1* to *C3*; Figure 4e; full results in Supplementary Table 1).

**Figure 3:**
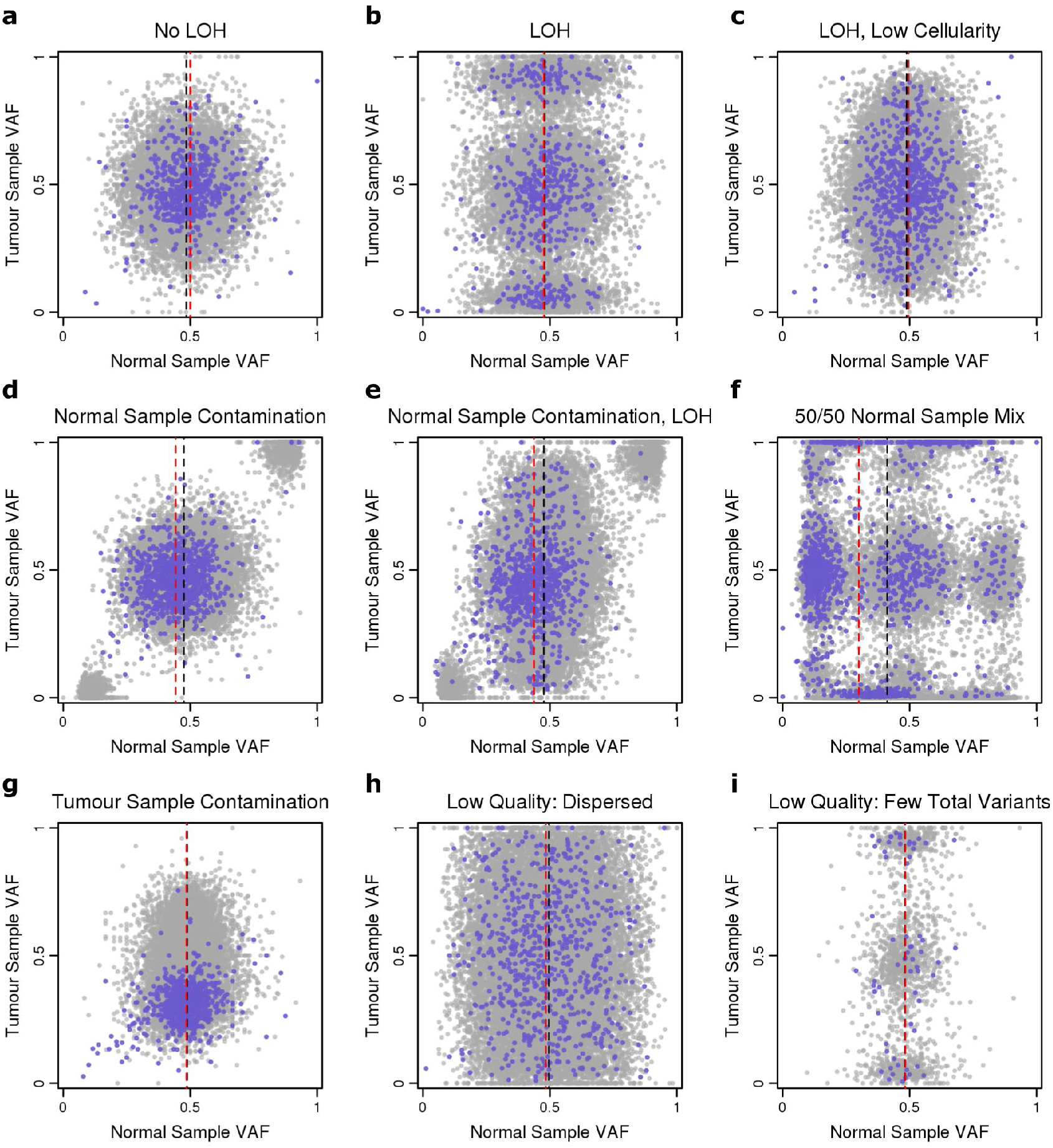
Example VAF distributions of germline heterozygous variants in paired tumour:normal samples. Rare variants = purple, common variants = grey. Dotted lines show the median normal sample VAF for rare (red) and common (black) variants. **a,** In tumour samples with no LOH, the VAF distributions in both samples cluster around 0.5. **bc,** Alleles that have undergone LOH in the tumour have a VAF close to 0 or 1. The degree of separation between non-LOH and LOH alleles is defined by the tumour cellularity (proportion of normal cells in the tumour biopsy) and the clonality of the LOH event (proportion of LOH cells in the tumour biopsy). **de,** Contaminated samples are characterised by a left-shift of rare variants, and clusters of variants with very low (<0.2) or very high (>0.8) VAF, caused by the mixing of different genotypes from two individuals - as seen in Supplementary Figure 4. **f,** A 50/50 mix of two normal samples splits the distribution into multiple clusters corresponding to the different genotype combinations - illustrated in Supplementary Figure 4a. **g,** Tumour sample specific contamination causes a down-shift of rare variants. Contaminating variants that do not appear in the normal sample are misinterpreted as somatic mutations (Table 1; Supplementary Figure 6). **hi,** Low sequencing quality can result in a highly dispersed distribution (**h**) or a very low total number of variants (**i**).

**Figure 4:**
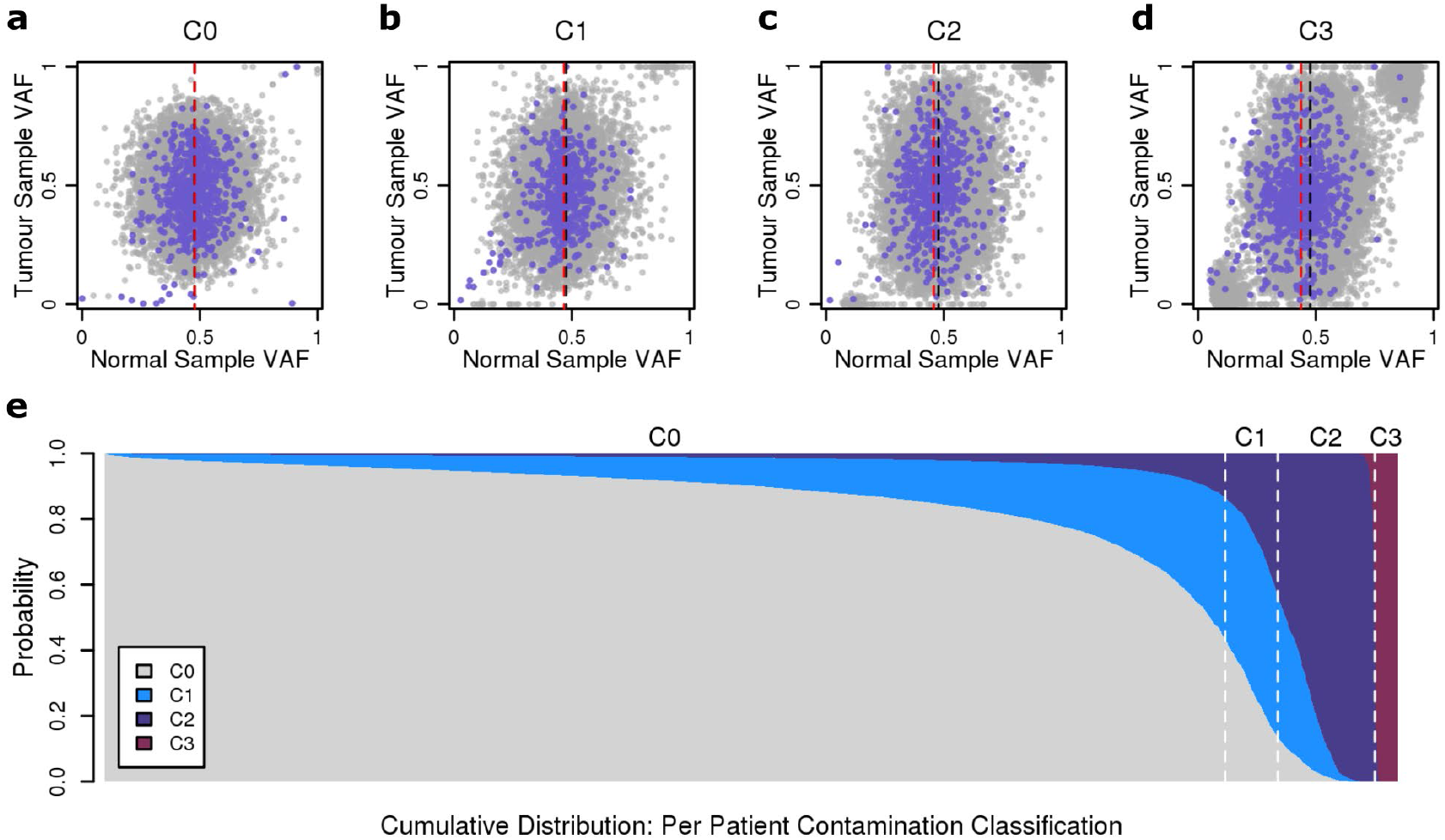
Evidence of substantial contamination in TCGA WXS data. **a-d,** Example VAF distributions of germline heterozygous variants in matched tumour:normal sample pairs from each contamination class. Rare variants = purple, common variants = grey. Dotted lines show the median normal sample VAF rare (red) and common (black) variants. **e,** Plot shows the calculated probability of each patient belonging to each contamination class, shown as a cumulative distribution. Probabilities are taken from the output of the ordinal logistic regression based classifier. Dotted white lines show the limits of the classes, ie: where the probability of belonging to that class is >0.5. Full results in Supplementary Table 1.

Further analysis of the contaminated samples identified a pair of patients, of the same cancer subtype, with an excessively high proportion of germline heterozygous variants in common; furthermore the shared variants formed distinct clusters in the tumour/normal VAF distribution (Supplementary Figure 4). Available metadata indicated they were likely processed in parallel (Supplementary Table 2) providing strong evidence of cross-contamination. Other corroborating metadata includes four patients with approximately 50% contamination in the normal sample, that were likely sequenced in parallel - indicative of mis-labelling of lanes during sequencing or mis-assignment of barcodes during downstream processing of the sequence data (Supplementary Figure 5, Supplementary Table 3).

**Table 1:**
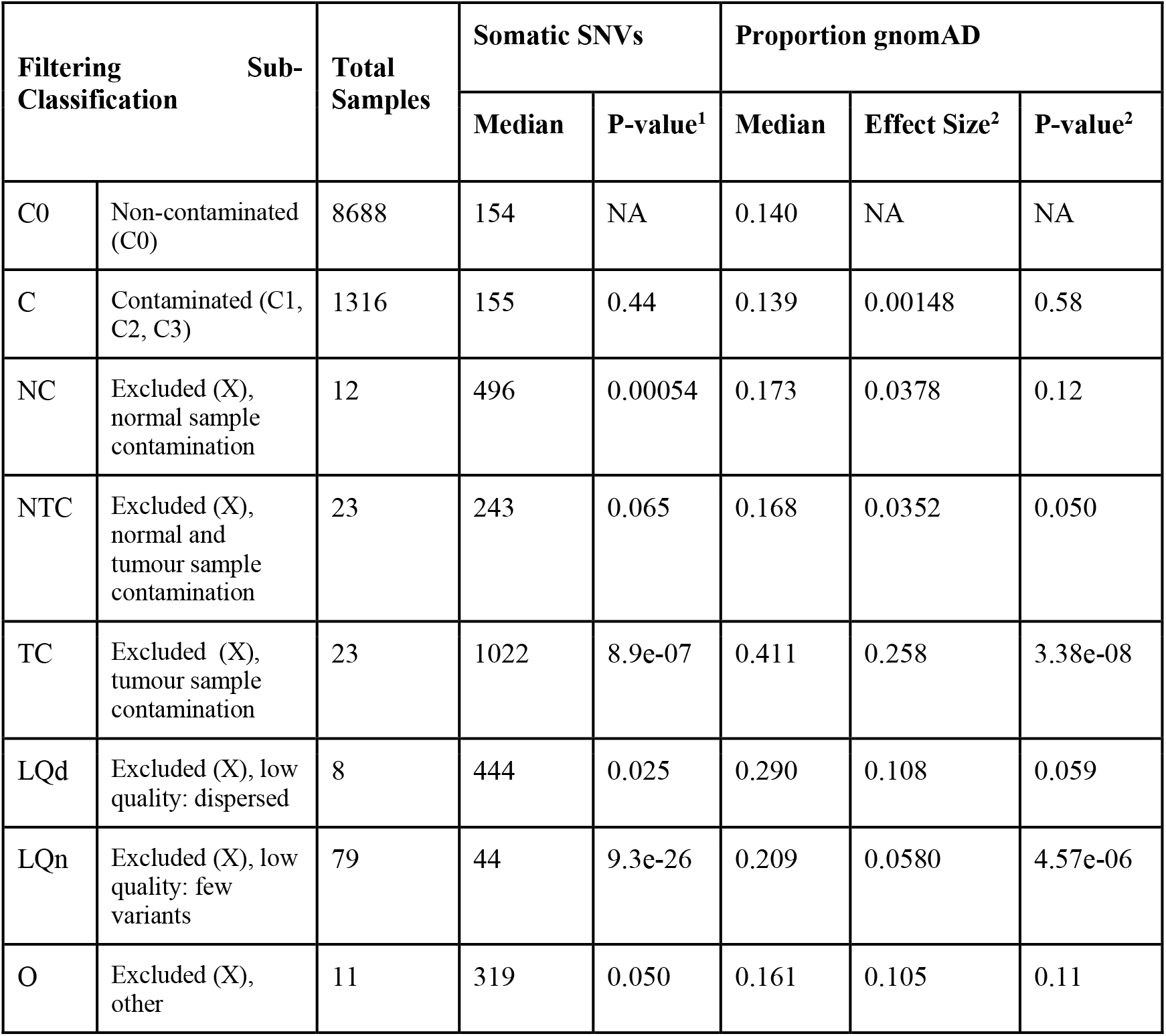
Influence of contamination and sequencing quality on somatic mutation data. Comparison of the total number of somatic SNVs in contaminated and abnormal samples, compared to non-contaminated samples (C0), and proportion of total somatic SNVs found in gnomAD. Excluded samples (X, n=185) were sub-classified based upon their reason for thresholding, samples identified as contaminated by logistic regression were combined into a single class (C). ^1^ Mann-Whitney U Test, versus C0 ^2^ Linear regression, response variable = proportion of somatic SNPs found in the gnomAD database, predictor variables = total somatic SNPs and filtering sub-classification.

Contamination and experimental error is not specific to TCGA and is a risk in all sequencing projects. To systematically identify such cases in tumour:normal pairs we have developed a software tool GenomeArtiFinder (https://git.ecdf.ed.ac.uk/taylor-lab/GenomeArtiFinder) allowing the easy application of the contamination classification and quantification methods developed here. Application to an in-progress study identified 4 problematic samples, independently confirmed using VerifyBamId^19^, out of 223 pairs of whole genome sequences (WGS). Application to a second in-progress study of 120 WGS sample pairs identified 3 sample swaps, confirmed using qSignature (available at: https://sourceforge.net/p/adamajava/wiki/qSignature/) and one highly contaminated sample, confirmed using VerifyBamId.

To assess the effect of contamination and other experimental errors on downstream analysis, we compared the total number of somatic single nucleotide variants (SNVs) within the different groups. The contaminated samples, identified by logistic regression (*C1*, *C2*, *C3*), were combined into a single class (*C*, n=1,316). The samples excluded in the first step (*X*, n=185) were sub-classified based upon their reason for thresholding, as: low quality with few total variants (*LQn*, n=79); low quality with highly dispersed VAFs (*LQd*, n=8); high normal sample specific contamination (*NC*, n=12); tumour sample specific contamination (*TC*, n=23); high contamination of both the tumour and normal sample (*NTC*, n=23) and other (*O*, n=11). *LQn* samples had significantly fewer somatic SNPs than non-contaminated (median total somatic SNPs 3.5 fold lower, Mann-Whitney U Test, p = 9.3e-26), whilst *NC* and *TC* had significantly more (Mann-Whitney U Test, p < 0.001; Table 1; Supplementary Figure 6a). We then calculated the proportion of each sample’s somatic SNVs that appeared in gnomAD - allowing us to infer the relative extent to which germline variants are being erroneously called as somatic SNVs. Using a linear regression model, we hence estimated the enrichment of somatic gnomAD variants within each group. All groups except *C* and *NC* had a significant increase in the proportion of somatic gnomAD variants compared to non-contaminated samples (p < 0.05), with the highest enrichment seen in *TC* samples (Supplementary Figure 6b; Table 1). For samples with evidence of tumour-sample contamination (*TC*, *NTC*), the increase in somatic gnomAD variants is likely due to mis-calling of contaminating germline variants in the tumour sample as somatic SNVs. In the low-quality samples (*LQn*, *LQd*), it seems likely that true-germline variants missed in the normal sample are erroneously called as somatic due to the lower stringency required for calling variants in tumour samples.

### Post-filtering analysis of biased allele retention

For the subsequent analysis, we excluded all thresholded samples (*X*) plus those classified as most severely contaminated (*C3*), leaving 9,602 patients in our total pan-cancer set. As the output of the classifier is quantitative, in instances where a single patient had multiple pairs of tumour:normal samples we retained only the least contaminated pair (Supplementary Figure 7). The artefactual variant and exome target capture kit bias filtering (as described above) was repeated with this patient set. Using the filtered patients and variants, we then repeated the pan-cancer LOH bias analysis. No variants appeared to be significantly preferentially retained during LOH (Supplementary Figure 8a; Bonferroni corrected p < 0.05). LOH bias analysis was performed individually for each cancer subtype with more than 75 samples (n = 30). After multiple-testing correction, no variants appeared significant in any of the analyses (Supplementary Figure 8b; Bonferroni corrected p < 0.05).

To estimate the minimum effect size our analysis has power to detect, we performed simulations based on the largest cohort (breast invasive carcinoma [BRCA]) and the cohort with the highest rate of LOH (ovarian serous cystadenocarcinoma [OV]) using the total number of patients (BRCA = 831, OV = 398, total patients = 9,602), and median frequency of LOH (BRCA = 0.31, OV = 0.45, total patients = 0.24; Supplementary Figure 8c-f). The results of the simulations indicated that for common variants (heterozygous frequency = 0.5), effect size odds ratios of 3.1 and 3.9 gave approximately 80% power to detect a robustly significant effect (Bonferroni corrected p-value < 0.05; BRCA and OV respectively). For less common variants (heterozygous frequency = 0.2), effect sizes of 7.8 and 16.0 were required. Although odds ratios for biased allele retention in established cancers are not directly comparable to odds ratios for cancer risk in the population, the maximum reported GWAS effect sizes in these cohorts was only 1.6 and 1.93 (EBI GWAS Catalog; BRCA and OV respectively). This suggests that with the data currently available we are underpowered to detect biased allele retention of common variants at an exome wide level of significance (Supplementary Figure 8e,f).

In a more targeted exploration, we tested for biased allele retention at previously reported GWAS significant variants in the matched cohort (e.g. lung cancer GWAS in lung cancer cohort). This potentially provides an orthogonal validation of the GWAS result and is an implicit test that the genetic effect is cancer cell autonomous, rather than for example manifesting as a cancer predisposition effect on the immune system. Cancer related SNPs from the EMBL-EBI GWAS Catalog (trait = cancer, EFO ID = EFO_0000311) were downloaded, and overlapped with our dataset. GWAS SNPs were then matched by trait to related TCGA cancer subtypes, leaving a final set of 172 SNPs, 118 with a reported GWAS OR. The LOH bias of GWAS SNPs from related traits was calculated and for variants with a reported GWAS OR, we used the observed number of LOH and non-LOH samples at the locus, and the GWAS OR to calculate the power of our analysis to detect a significant bias. Under these assumptions, none of the SNPs analysed here had sufficient power to detect a nominally significant LOH bias in this dataset (maximum power to detect a p-value < 0.05 = 35%; Supplementary Table 4; Supplementary Figure 9a). Despite this, 1 SNP from skin cutaneous melanoma (SKCM) was significant after multiple testing correction (OR = 0.10, Fisher’s exact test, Bonferroni adjusted p-value = 0.044), and a further 7 had an unadjusted p-value < 0.05 (5 lung squamous cell carcinoma [LUSC], 1 liver hepatocellular carcinoma [LIHC] and 1 rectum adenocarcinoma [READ]; Supplementary Table 4). In all cases, SNPs with a p < 0.05 and a reported GWAS risk allele (n = 6) were biased towards retention of the risk allele during LOH. Furthermore, the LOH bias ORs of the 118 SNPs with a reported GWAS OR were significantly correlated with the derived GWAS ORs (adjusted for direction of the reported risk allele; rho = 0.21, Spearman’s rank correlation, p-value = 0.020; Supplementary Figure 9b), indicating a significant trend towards retention of the risk allele. Together, this indicates that at least a subset of cancer-associated risk alleles are being selected by LOH during tumour development. Our ability to detect this effect despite low anticipated power in the analysis suggests that GWAS reported OR are underestimating the effect size in biased allele retention.

### LOH selects for disruptive germline variants in tumour suppressor genes and protein interaction pathways

Park *et al.^20^* recently reported selection of rare ‘potentially damaging’ germline variants during tumorigenesis via somatic LOH investigated using this same dataset. Analogously, we looked for evidence of selection of germline risk variants in genes from a curated catalogue of known cancer genes (COSMIC)^21^. Our analysis demonstrated a highly significant overall preference for retention of predicted loss-of-function mutations (annotated as ‘HIGH’ impact by Variant Effect Predictor [VEP^22^]; OR = 1.98, Fisher’s exact test, p = 6.7e-09; Figure 6a) and known pathogenic mutations (‘Pathogenic’ and ‘Likely Pathogenic’ annotations from ClinVar^23^; OR = 3.11, Fisher’s exact test, p = 9.2e-12; Figure 6b) in tumour suppressor genes with a recessive phenotype. In total, 418 out of 1,175 individuals (35.57%) with a heterozygous predicted damaging germline variant in a recessive tumour suppressor gene underwent LOH at the locus, and of these 284 (67.94%) retained the damaging allele.

We subsequently performed LOH bias tests individually for all COSMIC genes with at least 1 heterozygous predicted damaging germline variant. ATM, BRCA1, BRCA2 and SDHB were significantly biased towards retention of the damaging allele (false discovery rate [fdr] < 0.05; Figure 6c; Supplementary Table 4). Preferential retention of damaging ATM, BRCA1 and BRCA2 germline variants has previously been noted in the TCGA samples^20^. The LOH analysis was then repeated for all protein coding genes. Although no further genes appeared significant after multiple testing correction, PADI3 showed evidence of purifying oncogenic selection, with a nominally significant bias towards retention of the non-damaging allele (OR = 0.54, Fisher’s exact test, p = 0.0010; Figure 6d). This result indicates that functionally active PADI3 may be essential within a subset of the tumours, a result that is supported by previous *in vivo* studies of PADI enzymes^24^.

Finally, we looked for evidence of selection of predicted damaging germline variants in protein interaction pathways from the pathway database, Reactome^25^. Out of 1,897 pathways tested, 25 had a significant bias towards retention of the damaging allele after LOH (Bonferroni corrected p-value < 0.05; Figure 6e,f; Table 2). These pathways were split between five biological processes: DNA repair (n=14), gene expression (n=5), cell cycle (n=4), metabolism of proteins (n=1) and reproduction (n=1). In total, the significant pathways included 232 unique genes, 12 of which, when removed from the analysis, cause at least one pathway to drop below the threshold for significance, demonstrating a significant contribution towards the overall burden of that pathway (Figure 6g; ATM, BRCA1, BRCA2, BRIP1, HIST1H4B, HIST1H4H, MCM2, MRE11, NBN, TP53, UBA52 and WRN). At least one of: ATM, BRCA1 or BRCA2 appeared in every significant pathway, indicating that these individually significant genes were contributing the majority of the signal.

**Table 2:** Reactome Pathways with significant preferential retention of predicted damaging germline variants during loss-of-heterozygosity. LOH bias test of predicted damaging germline variants was performed for all Reactome protein interaction pathways containing at least one gene with a heterozygous predicted damaging germline variant (n=1,897). Table shows all significant pathways: Bonferroni corrected p-value < 0.05. ^1^ Fisher’s exact test, rare versus common allele retention of damaging versus benign variants ^2^ Bonferroni adjusted p-value

| Biological Process | Pathway Identifier | Pathway Name | ID | Total Genes | OR<sup>1</sup> | p-value<sup>1</sup> | Adj p-value<sup>2</sup> |
| --- | --- | --- | --- | --- | --- | --- | --- |
| Repair | R-HSA-5693532 | DNA Double-Strand Break Repair | D2 | 94 | 1.71 | 4.0e-08 | 7.2e-05 |
|  | R-HSA-5693606 | DNA Double Strand Break Response | D3.a | 37 | 2.77 | 8.1e-11 | 1.5e-07 |
|  | R-HSA-5693538 | Homology Directed Repair | D3.b | 79 | 1.73 | 1.4e-07 | 0.00025 |
|  | R-HSA-5693571 | Nonhomologous End-Joining (NHEJ) | D3.c | 38 | 2.55 | 1.9e-09 | 3.6e-06 |
|  | R-HSA-5693565 | Recruitment and ATM-mediated phosphorylation of repair and signaling proteins at DNA double strand breaks | D4.a | 37 | 2.77 | 8.1e-11 | 1.5e-07 |
|  | R-HSA-5693567 | HDR through Homologous Recombination (HRR) or Single Strand Annealing (SSA) | D4.b | 74 | 1.83 | 1.6e-08 | 2.9e-05 |
|  | R-HSA-5685942 | HDR through Homologous Recombination (HRR) | D5.a | 47 | 2.44 | 8.8e-13 | 1.6e-09 |
|  | R-HSA-5685938 | HDR through Single Strand Annealing (SSA) | D5.b | 26 | 1.94 | 1.6e-05 | 0.029 |
|  | R-HSA-5693607 | Processing of DNA double-strand break ends | D5.c | 51 | 1.99 | 1.6e-06 | 0.0030 |
|  | R-HSA-5693579 | Homologous DNA Pairing and Strand Exchange | D6.a | 33 | 2.98 | 5.4e-15 | 1.0e-11 |
|  | R-HSA-5693537 | Resolution of D-Loop Structures | D6.b | 29 | 3.18 | 2.6e-16 | 4.8e-13 |
|  | R-HSA-5693616 | Presynaptic phase of homologous DNA pairing and strand exchange | D7.a | 30 | 2.93 | 9.1e-14 | 1.7e-10 |
|  | R-HSA-5693568 | Resolution of D-loop Structures through Holliday Junction Intermediates | D7.b | 28 | 3.24 | 1.2e-16 | 2.2e-13 |
|  | R-HSA-5693554 | Resolution of D-loop Structures through Synthesis-Dependent Strand Annealing (SDSA) | D7.c | 24 | 3.47 | 2.9e-17 | 5.4e-14 |
| Gene Expression | R-HSA-8953750 | Transcriptional Regulation by E2F6 | G4.a | 11 | 3.77 | 3.1e-07 | 0.00057 |
|  | R-HSA-6796648 | TP53 Regulates Transcription of DNA Repair Genes | G5.c | 35 | 2.30 | 1.1e-08 | 2.1e-05 |
|  | R-HSA-6804760 | Regulation of TP53 Activity through Methylation | G6.a | 9 | 3.27 | 2.1e-05 | 0.038 |
|  | R-HSA-6804756 | Regulation of TP53 Activity through Phosphorylation | G6.b | 42 | 2.05 | 1.3e-07 | 0.00023 |
|  | R-HSA-6803207 | TP53 Regulates Transcription of Caspase Activators and Caspases | G6.c | 8 | 3.34 | 1.5e-05 | 0.028 |
| Cell Cycle | R-HSA-1500620 | Meiosis | C2.b | 59 | 1.84 | 9.0e-09 | 1.6e-05 |
|  | R-HSA-69481 | G2/M Checkpoints | C3.a | 75 | 1.67 | 1.5e-05 | 0.027 |
|  | R-HSA-912446 | Meiotic recombination | C3.b | 40 | 2.23 | 5.1e-11 | 9.5e-08 |
|  | R-HSA-69473 | G2/M DNA damage checkpoint | C4 | 51 | 2.00 | 6.0e-07 | 0.0011 |
| Metabolism of Proteins | R-HSA-5689901 | Metalloprotease DUBs | M4 | 17 | 4.13 | 5.1e-08 | 9.3e-05 |
| Reproduction | R-HSA-1474165 | Reproduction | R1 | 78 | 1.63 | 4.7e-07 | 0.00086 |

Overall, our analysis found strong evidence for selection of predicted damaging germline variants in tumour suppressor genes during LOH as a common mechanism of cancer evolution. Furthermore, we identified proteins and pathways involved in DNA repair as the most significant targets.

## Discussion

LOH as a mechanism of oncogenic selection has been a key tenet of cancer etiology since 1971^26^. By performing a targeted analysis of potentially deleterious germline variation in known cancer associated genes, we identified LOH as the ‘second-hit’ in 24.17% of patients with a heterozygous damaging variant in a tumour suppressor genes (n = 284; 2.96% of 9,602 total patients analysed). Analysis of individual genes found this signal to be dominated by mutations in: ATM, BRCA1, BRCA2 and SDHB (Figure 5c). As inactivation of ATM, BRCA1 and BRCA2 are all associated with increased genome instability and consequent LOH^27–29^, this indicates a cyclical mechanism of oncogenesis, in which a heterozygous loss-of-function mutation reduces the overall efficiency of double-strand break repair (DSBR), thereby increasing the likelihood of a LOH event targeting the WT copy. This conclusion is further supported by studies demonstrating that whilst BRCA1 and BRCA2 heterozygous knockouts are phenotypically normal under most conditions, a phenotype of ‘conditional haploinsufficiency’ and consequent genomic instability can be induced by exposure to different endogenous and exogenous stressors^30–33^. The identification of significant preferential retention of predicted damaging germline variants in DSBR pathways (Figure 5e), including significant contributions from other DSBR proteins such as: MRE11, WRN and NBN (Figure 5g) implies that genomic instability driven by conditional haploinsufficiency of DSBR proteins may be a common cancer evolution pathway. By studying the effects of different stressors in cells with heterozygous DSBR loss-of-function mutations, we will gain insight into the specific combination of genetic and environmental factors that drive the ‘second hit’ in cancer evolution.

**Figure 5:**
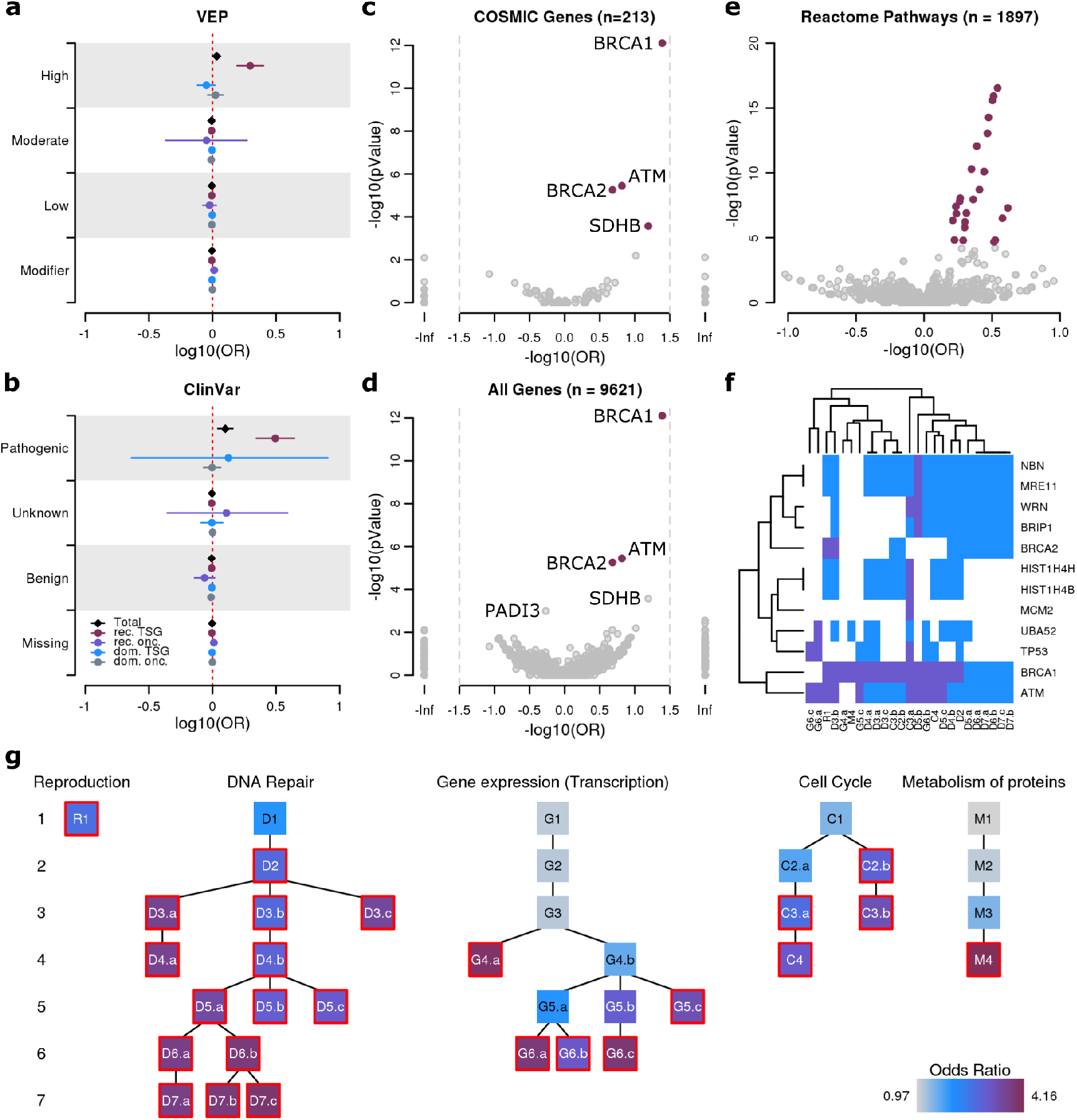
Selection for predicted damaging germline variants via LOH during tumour development. **ab,** LOH bias for COSMIC genes. Genes were grouped by annotations in COSMIC and either (**a**) VEP or (**b**) ClinVar. Horizontal lines show the 95% confidence interval. **c,** LOH bias of predicted damaging germline variants in COSMIC genes. Purple = fdr < 0.05. Full results in Supplementary Table 5. **d,** LOH bias of predicted damaging germline variants in genes with at least 1 predicted damaging germline variant. **e,** LOH bias of predicted damaging germline variants in Reactome pathways. Purple = p-value < 0.05. In **a-e**, log10(OR) <0 = reference allele bias, >0 = alternative allele bias. **f,** Pathway membership of genes found to contribute to the overall burden of at least one significant pathway. Blue indicates pathway membership, purple indicates that when removed from the analysis the pathway drops below the threshold for significance. **g,** Hierarchical network diagram of Reactome pathways with a significant LOH bias towards predicted damaging germline variants (Bonferonni corrected p-value < 0.05). Each node represents a biological pathway, statistically significant pathways have a red border. Branches link sub-processes within larger pathways, with the most broadly defined biological processes at the top of the plot. Non-significant branches within each network are excluded. Details of each significant pathway are shown in Table 2.

Our results show that biased LOH selects for pathogenic or loss-of-function variants in recessive tumour suppressor genes (Figure 5a,b), demonstrating the dysregulation of normal cellular function as a key step in the acquisition of oncogenicity. In contrast, PADI3 (peptidylarginine deiminase 3) showed nominally significant purifying selection against predicted damaging germline variants (OR = 0.54, t-test, unadjusted p = 0.0010; Figure 6d), indicating that functionally active PADI3 may be necessary for a subset of tumours, a result that is supported by previous *in vivo* studies of PADI enzymes^24^. Due to the high proportion of BRCA and OV patients in the TCGA cohort (831 and 398, respectively; 12.80% of total patients), these results are predominantly biased towards breast and ovarian cancer associated genes - by repeating this analysis within different cohorts of patients, it may be possible to identify functionally essential proteins from specific cancer subtypes, revealing novel therapeutic targets.

The majority of LOH events do not involve loss-of-function or pathogenic variants, and hence their phenotypic impact is still unknown^5^. It is possible that many common LOH events are passenger mutations with no functional influence on cancer progression, or alternatively, it is perhaps more likely that due to the limitations of the data currently available, we are unable to detect the selective effect of common, small effect variants. In this study we identified a significant correlation between the GWAS OR and LOH OR of previously reported GWAS variants in matched cancer subtypes (Supplementary Figure 9b), demonstrating a pan-cancer trend towards retention of cancer-associated variants during LOH. Taken with previous analyses of biased LOH at putative risk variants^5–7^, this result indicates that small-effect cancer associated variants are likely influencing cancer evolution, albeit to an extent that we are currently underpowered to robustly detect per-gene.

TCGA is one of the most widely used resources in cancer genomics, with more than 2,000 citations of the original 2013 flagship publication^8^ and over 3,500 papers in pubmed containing the keyword ‘TCGA’. Consequently, our identification and quantification of substantial contamination and experimental error in 382 pairs of samples (185 = *X*; 197 = *C3*), and mild to moderate contamination in a further 1,296 (*C1* = 456; *C2* = 840) has important implications. For example, our discovery of a significant increase in total somatic SNP burden in contaminated samples (Supplementary Figure 6a) illustrates how genetic material from other sources can be mis-interpreted as somatic mutations, potentially leading to incorrect conclusions. Batch effects have previously been reported across TCGA samples^9,12^ but these studies have not considered sample contamination as a systematic confounder. We provide our per-sample contamination estimates as a resource to allow other researchers working with TCGA to choose appropriate filters for their analysis (Supplementary Table 1). Furthermore, our results and methodology provide a broadly applicable framework that can be used to profile contamination, low quality data and other technical issues using germline variant data from matched tumour:normal samples. To enable the general application of these quality control metrics to both user-generated and public data we make the GenomeArtiFinder software package available (https://git.ecdf.ed.ac.uk/taylor-lab/GenomeArtiFinder).

As we’ve demonstrated, the primary limiting factor in genomics studies is power (Supplementary Figure 8c-d) - consequently, the genomics field is becoming dominated by large collaborative sequencing projects, relying on data generated in multiple batches, from multiple centres, often over multiple years. Many projects - including TCGA and PanCancer Analysis of Whole Genomes (PCAWG)^34^ - use standardised bioinformatic pipelines to eliminate technical variation in the downstream analysis, and although important - this does nothing to account for experimental variation. The confounding impact of experimental batch effects in high-throughput data and their propensity to lead to false conclusions has been well-documented^35,36^, and yet they often remain unaccounted for. Many batch effects can be overcome with careful experimental design - for example: ensuring comparative groups (eg: test samples and controls) are processed in parallel to avoid confounding technical variation with biological difference; and using standardised reagents and protocols to minimise experimental variation. Additionally, experimental metadata - such as sample processing and sequencing groups - should be made readily available to researchers, so that the potential confounding impact can be properly profiled, and where necessary accounted for.

Measures of biased allele retention efficiently re-discover loci known to harbour cancer predisposing germline variants in the human population, and implicate new candidates such as MCM2. Pathway based analyses of these rare deleterious variants shows their importance to many families, and points to a genome instability ratchet that biases heterozygous carriers of deleterious mutations to undergoing LOH providing an opportunity to develop a full-blown genome instability phenotype. This ratchet effect may also explain why GWAS odds ratios seem to under-estimate the effect size of damaging variants when viewed from the perspective of biased allele retention. The success of GWAS validation suggests that the biased allele retention approach will be informative at the genome-wide scale as larger datasets of cancer genome sequencing are acquired.

## Materials and Methods

### Availability of data and materials

Details of all software and packages used in the analysis are in Supplementary Table 6.

All analyses were performed using the Genomic Data Common (GDC) data harmonization and generation pipeline GRCh38 reference sequence (GRCh38.d1.vd1.fa, available from: https://gdc.cancer.gov/about-data/data-harmonization-and-generation/gdc-ference-files). Aligned whole exome sequence (WXS) reads were downloaded as BAM files from the TCGA and TARGET projects using the GDC data portal. Somatic variant calls generated from matched tumour:normal pairs were downloaded from the GDC data portal as VCFs. Details of BAM and VCF pre-processing are available from: https://gdc.cancer.gov/documentation.

Variant calling and LOH/SCNA analysis were limited to exonic regions using a genomic region file, adapted from ExAC (exome_calling_regions.v1.interval_list, available from: ftp://ftp.broadinstitute.org/pub/ExAC_release/release0.3.1/resources/). Firstly, intervals were lifted over from hg19 to hg38 using Picard (v2.6) and LiftOver resources from UCSC Genome Browser. Secondly, genomic intervals of low-complexity regions (LCR) and segmental duplication (SegDup) regions were downloaded from gnomAD (LCR.interval_list and mm-2-merged.bed.gz, respectively, both available from gs://gnomad-public/intervals/ using the Python application gsutil), and lifted over to hg38. LCR and SegDup regions were then subtracted from the exome target region file using BEDTools (v2.25.0)^37^.

To produce a consensus file of genomic regions common to all exome target capture kits used in TCGA, BED files corresponding to the target regions of each of the kits were queried from TCGA metadata. BED files of additional probes and target regions were excluded, as well as Gapfiller_7m (all patients sequenced using this kit failed filtering steps) and SureSelect_50Mb (only used to sequence 5 patients); consequently BED files from eight exome target capture kits were downloaded (Supplementary Table 7). Files were lifted over to hg38 then intersected using BEDTools to produce a consensus file of genomic regions common to all eight kits. Finally, LCR and SegDup regions were then subtracted from the exome target region file as described previously.

GnomAD VCFs from WXS and WGS (gnomad.exomes.r2.1.sites.vcf.bgz and gnomad.genomes.r2.1.sites.vcf.bgz, available for download using gsutil (v4.28) from gs://gnomad-public/release/2.1.1/vcf/) were downloaded then lifted-over to hg38 using Picard (v2.9.4). Multi-variant positions were split using a custom perl script, to ensure correct assignment of annotations to alternative alleles. Normalisation and further processing was performed using bcftools (v1.3.1).

### Sample Selection

For each patient, one pair of tumour:normal samples were selected for analysis, ensuring that both samples were prepared for sequencing using the same exome target capture kit and where possible were sequenced in the same experiment. For patients with multiple pairs of samples, the most recently sequenced pair was chosen. Out of 10,316 patients, 9,905 had at least one pair of tumour:normal samples that passed our criteria and underwent successful LOH prediction. Following contamination classification, only sample pairs classified as ‘C0’, ‘C1’, and ‘C2’ were used in the analysis. For patients with multiple pairs of samples, the pair with the lowest contamination score was selected (examples in Supplementary Figure 7). In total, 9,602 patients were included in the final LOH bias analysis.

### Germline Variant Calling and Annotation

Germline variants were called from normal BAM files individually using Strelka2 (v2.8.3)^13^ and in batches of approximately 200 samples using the GATK HaplotypeCaller (v3.8) best practices workflow^14,38^. The output VCFs from Strelka2 and GATK HaplotypeCaller were then overlapped and filtered to keep only variants that were called by both callers and passed all quality control measures. Variant annotation was performed using Variant Effect Predictor (VEP)^22^ using the v88 GRCh38 cache (homo_sapiens_vep_88_GRCh38.tar.gz, available from ftp://ftp.ensembl.org/pub/release-88/variation/VEP/).

### Loss of Heterozygosity and Somatic Copy Number Alteration Prediction

CloneCNA (v2.0) was used to predict changes in copy number and heterozygosity, and additionally estimate tumour cellularity and clonality of mutations^15^.

### Exome-Wide Preferential Allelic Imbalance Detection

Allele specific read counts for germline heterozygous SNPs were generated individually for all pairs of tumour:normal BAM files using ExomeSeqMiner (included with the CloneCNA software package). SNPs with VAF <0.2 or >0.8 or read depth <10 in the normal sample were removed from the analysis. Loci were then overlapped with segmental LOH/SCNA predictions from CloneCNA to give a per-variant LOH prediction. For each variant, the allelic count odds ratio (OR_AC_) of alternative versus reference reads in tumour versus normal was calculated to predict which allele had been retained in the tumour. At every loci with at least 50 heterozygous individuals within the test group, a Fisher’s exact test was performed comparing counts of test group samples that had undergone LOH at the loci (CloneCNA predictions: HEMD [hemizygous deletion], NLOH [copy-neutral loss-of-heterozygosity] or ALOH [copy-amplification loss-of-heterozygosity]) and had retained the alternative or reference allele (OR_AC_ >1 or OR_AC_ <1, respectively), versus counts of all patient samples that had not undergone LOH or any other segmental mutation at the loci (CloneCNA prediction: NHET [copy-neutral heterozygous)]). We expect that the OR_AC_ of NHET samples should be equally distributed around 1, therefore by using the NHET samples as a control group, we can control for overall bias in the distribution of OR_AC_ at a given loci. Exome-wide LOH bias tests were performed for the pan-cancer dataset (all versus all), and on a cancer subtype specific basis (test group versus all).

### Cancer GWAS Variants

Using the EMBL-EBI GWAS Catalog, all previously reported cancer-associated variants were downloaded (trait = cancer, EFO ID = EFO_0000311, 5,552 associations from 580 studies). After processing, genome information was extracted for 4,723 unique SNPs. SNPs were intersected with our dataset, leaving 217 cancer associated exome variants. For each cancer subtype, associated traits were queried with related keywords to extract cancer subtype specific SNP associations. LOH bias was calculated for each SNP as described above.

### COSMIC: Preferential Allelic Imbalance Detection

To test for preferential selection of mutations in known cancer genes, tier 1 genes from COSMIC (Catalog of Somatic Mutations in Cancer; cancer_gene_census.csv, available from: https://cancer.sanger.ac.uk/cosmic/download) were grouped by role (oncogene versus tumour suppressor) and molecular genetics (recessive versus dominant). Germline variants within each of these gene groups were then collected and cross-referenced with their VEP annotations and ClinVar^23^ predictions of pathogenicity (clinvar_20190325.vcf.gz, available from: ftp://ftp.ncbi.nlm.nih.gov/pub/clinvar/vcf_GRCh38/), in addition to their LOH status. For each gene group, and each VEP annotation category (‘HIGH’, ‘MODERATE’, ‘LOW’, ‘MODIFIER’), and ClinVar prediction (‘Pathogenic’, ‘Benign’, ‘Unknown’, ‘Missing’; ClinVar categories were collapsed as shown in Supplementary Table 8) a Fisher’s exact test was performed comparing retention of the reference versus alternative alleles in samples that underwent LOH versus those that didn’t, as described above.

### Gene and Gene Pathway: Preferential Allelic Imbalance Detection

To test for preferential selection of potentially damaging mutations, germline heterozygous variants in all genes were collected and cross-referenced with their VEP annotations and ClinVar predictions of pathogenicity. On a gene-by-gene basis, reference versus alternative allele retention after LOH in samples with at least one damaging mutation (annotated as ‘HIGH’ impact by VEP, with a population allele frequency of <0.001 (gnomAD); or annotated as ‘Pathogenic’ in ClinVar) was compared to reference versus alternative allele retention after LOH samples with benign mutations (annotated as ‘LOW’ impact or ‘MODIFIER’ by VEP, with a population allele frequency of >0.05; or annotated as ‘Benign’ in ClinVar) using a Fisher’s exact test.

To test for enrichment in specific gene interaction pathways, a list of all genes with at least one predicted damaging germline variant were entered into Reactome^25^. Genes were then grouped into pathways identified by Reactome, and Fisher’s exact tests were performed combining counts from all genes within each pathway.

### Mapping And Sequencing Artefacts

To quantify the ‘reliability’ of all common variants (heterozygous frequency >1%), we used a binomial test to calculate the probability of sampling the observed VAF at the observed read depth based upon the expected reference allele bias (0.531^39^). Read depths were first normalised as follows: *(variant read count / sample median read depth) * population median read depth.* A binomial test was then performed for every common variant in every heterozygous normal sample. For each variant, we then calculated the proportion of the 99% confidence intervals given by the binomial test that overlapped with the expected - to give an overall measure of reliability at that locus. Finally, we compared the proportion overlapping the expected across all common variants in the dataset, and selected a threshold at the apex of the distribution (0.88, Figure 1f).

### Exome Target Capture Kit Bias

To quantify the extent of intra-kit allele-sharing, we performed pairwise correlations between randomly selected patients from the seven most common kits used in TCGA. Firstly, 50 patients were randomly selected from each kit. Secondly, the heterozygosity of each patient was determined at 5,000 randomly selected variants appearing in genomic regions common to all exome target capture kits and heterozygous in at least 50 of the sampled patients. Thirdly, heterozygosity was correlated between each pair of patients, to give a measure of allele sharing. This analysis was permuted 12 times using different randomly selected sets of patients and variants. Correlation coefficients were then compared between all kits.

To identify kit-specific enrichment of variants, we used linear regression to calculate the effect of each kit on the observed population frequency of heterozygotes (hetFreq), compared to the expected (estimated from the gnomAD non-Finnish European genome allele frequency [NFE_AF], using Hardy-Weinberg equilibrium). HetFreq was estimated from White/European individuals only, to account for population stratified variants. Patients were grouped by kit and by cancer subtype - to control for the possibility of cancer segregating germline variants that may be over-represented in specific kit groups. To account for variation in sample size, the linear regression was weighted by the number of individuals within each kit/cancer-subtype group.

### Manually Excluded Variants

Whilst investigating abnormal patient VAF distributions, we identified 25 patient samples (24 prostate adenocarcinoma [PRAD] and 1 kidney renal papillary cell carcinoma [KIRP]) with multiple rare variants with low VAF in both the tumour and normal samples. After cross-referencing these variants, we found that although they were rare across the total dataset, they were enriched within this subset of samples. We consequently compared their observed hetFreq within these 25 patients to their expected hetFreq (estimated from the gnomAD population allele frequency using Hardy-Weinberg equilibrium) and filtered any variant that was more than fivefold enriched, removing 170 variants in total. All analyses were performed having pre-filtered these variants from all samples.

### Thresholding of Contaminated, Low Quality and Abnormal Samples

Firstly, tumour/normal VAF distributions were split into grids of 25 evenly sized squares, and the number of rare and common variants appearing in each square counted, giving a vectorised representation of the total distribution. Secondly the median VAF was calculated for rare and common variants in both the tumour and normal samples - giving an estimate of any strong shifts in the overall distribution, and allowing identification of samples with lower VAF in rare variants compared to common - a characteristic of contaminated samples. The standard deviation of VAF in the normal sample was also calculated, to identify samples with high VAF dispersion - indicative of low quality sequence data. Finally, the proportion of rare and common variants above and below the midpoint of the central distribution was calculated, to give another representation of any shifts in the distribution of variants in the tumour sample - often the result of tumour-sample specific contamination. Individual examples from across the distribution of the different variables were investigated, and thresholds were consequently placed to remove individuals with abnormal tumour/normal VAF distributions.

### Sample Specific Quantification of Contamination

To construct a quantitative contamination classifier, a combination of 22 metrics that best captured the features of the tumour/normal VAF distributions contributable to contamination were chosen (variant counts in the outer regions of the tumour/normal VAF distribution [normal sample VAF <0.2 or >0.8] and median normal sample VAF of rare and common variants). Using the chosen metrics, we calculated the Mahalanobis distance of all samples from the total distribution. After splitting the ranked distribution of Mahalanobis distances into 6 groups, 100 patients were randomly selected from within each group (600 patients total) to generate a training set, equally representing the complete scale of contamination seen in our dataset. Patients were then manually classified as non-contaminated (C0) or contaminated (C1, C2, C3) using a sliding scale of severity, with ‘C3’ representing the most severe contamination. An ordinal logistic regression was performed on these samples, using the 22 metrics as predictor variables, and contamination group as the outcome variable. The output was then used to systematically classify the whole dataset.

### Somatic Variant Analysis

For each tumour:normal sample pair, somatic variant VCFs from four different somatic variant callers (MuSE, MuTect2, VarScan2, SomaticSniper) were downloaded from GDC. VCFs were intersected using GATK CombineVariants, and only variants passing all filters in at least two of the variant calling pipelines were kept.

To assess the extent of contaminating germline variants in the tumour samples, somatic variants from each tumour sample were overlapped with gnomAD exome variants, and the proportion of total SNPs appearing in the gnomAD database was calculated.

## Funding

This work was supported by the UK Medical Research Council, MRC Human Genetics Unit core funding programme grant MC_UU_00007/16. JL is supported by a PhD fellowship under the MRC Human Genetics Unit core funding programme.

## Author information

JL and MST designed the study. JL implemented and performed the analyses. RSY and AMM advised on analysis. JL and MST wrote the manuscript. All authors approved the final manuscript.

## Supplementary Figures

**Supplementary Figure 1:**
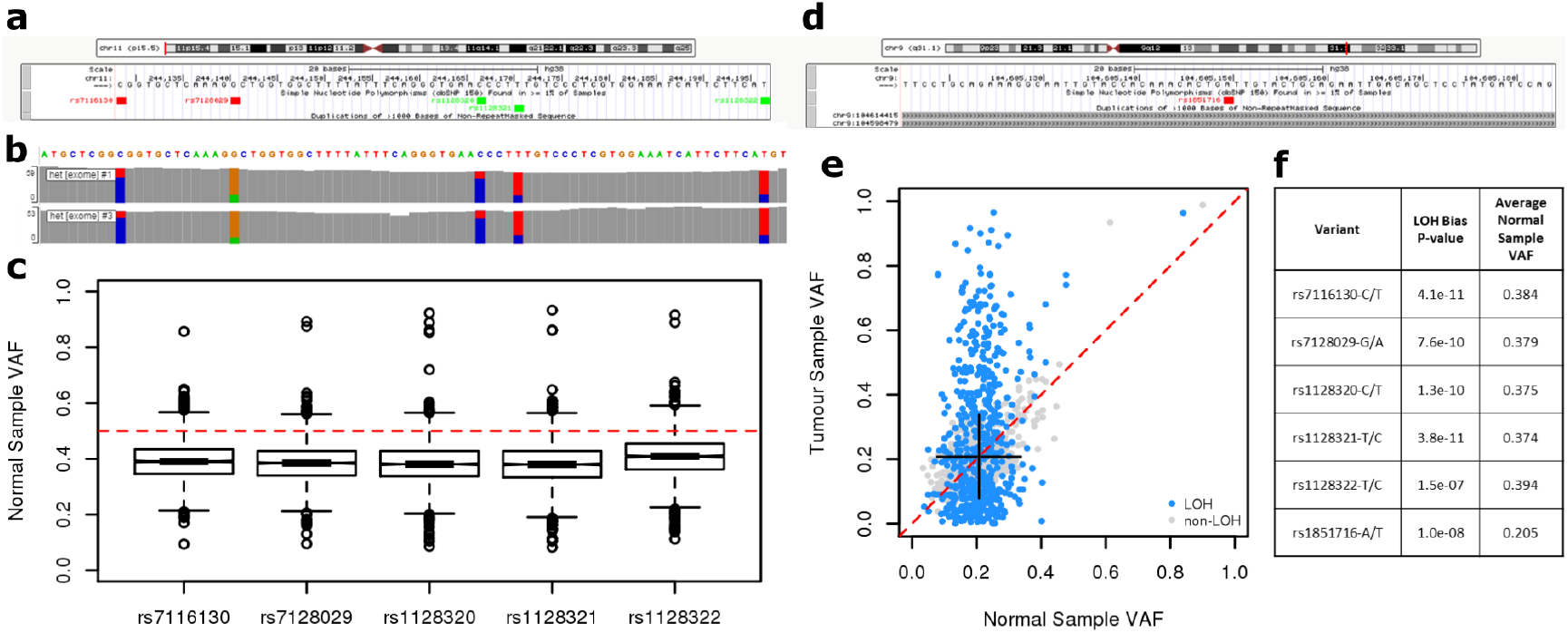
Regions of high-sequence identity and fixed haplotype blocks affect read alignment. **a,** Screenshot from the UCSC genome browser of chr11:244129-244197. **b,** Screenshot from gnomAD interactive IGV.js showing read alignments of whole exome sequencing data (WES) from normal samples. Coloured bars indicate proportion of reads containing the indicated base at that position. **c,** Boxplots of normal sample variant allele frequency (VAF) of the indicated variants from heterozygous individuals. Red dotted line indicates the expected VAF for a heterozygote (0.5). **d,** Screenshot from the UCSC genome browser of chr9:104605116-104605184. **e,** VAF of rs1851716-A/T in matched tumour:normal samples from germline heterozygous individuals. Black lines show the limits of 2*sd+mean of normal sample VAF. Red dotted line: x=y. **f,** Table of variants included in this figure.

**Supplementary Figure 2:**
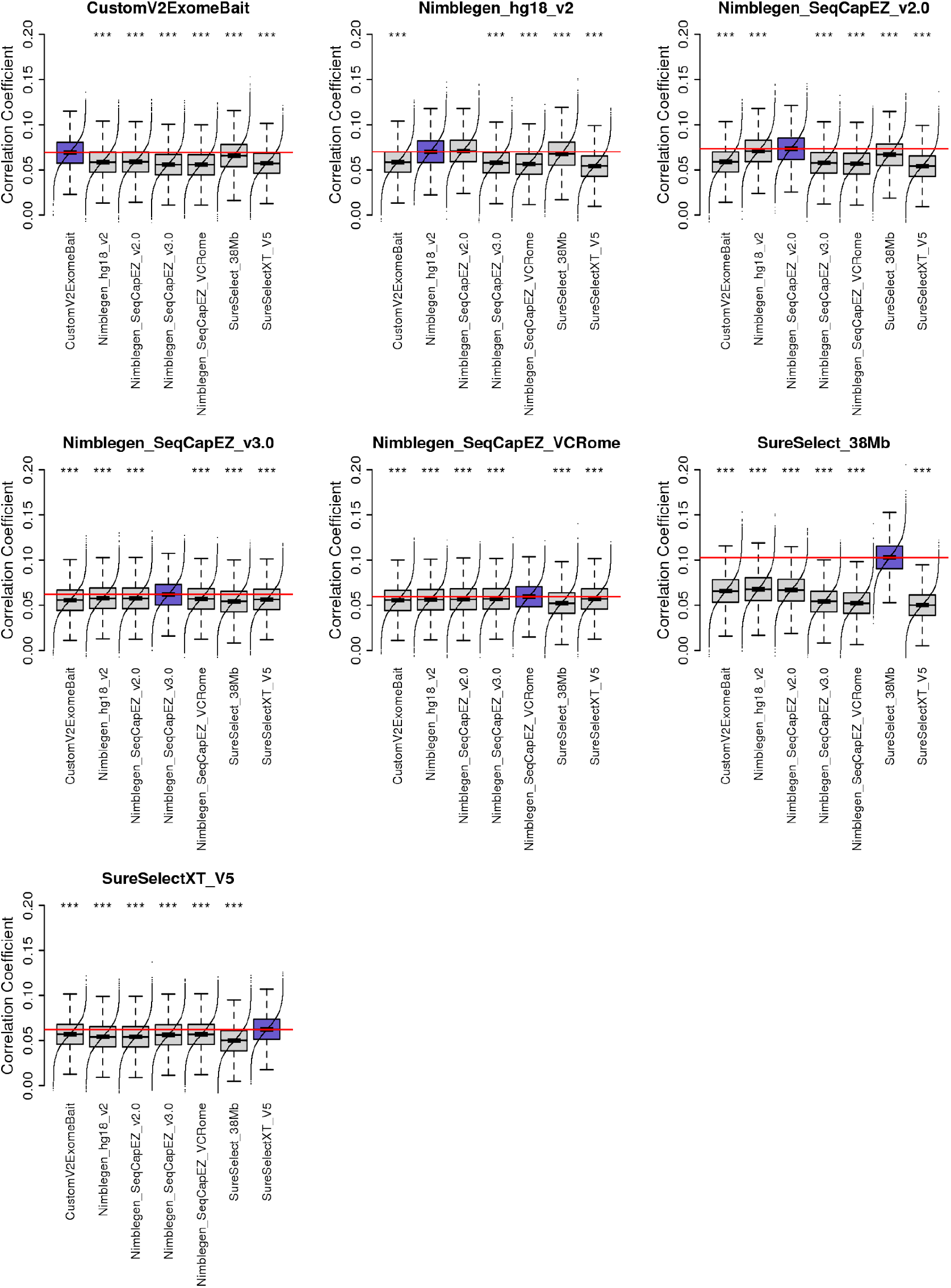
Intra-kit correlations. Boxplot plots show correlation coefficients of pairs of patients sequenced by the indicated kits. Analysis was permuted 12 times, for 50 randomly selected patients from each kit, comparing heterozygosity at 50,000 common variants. The coloured boxplot in each plot shows the intra-kit comparison. The red horizontal line shows the median of correlation coefficient of the intra-kit comparison. *** = p-value < 0.001, t-test compared to the intra-kit comparison.

**Supplementary Figure 3:**
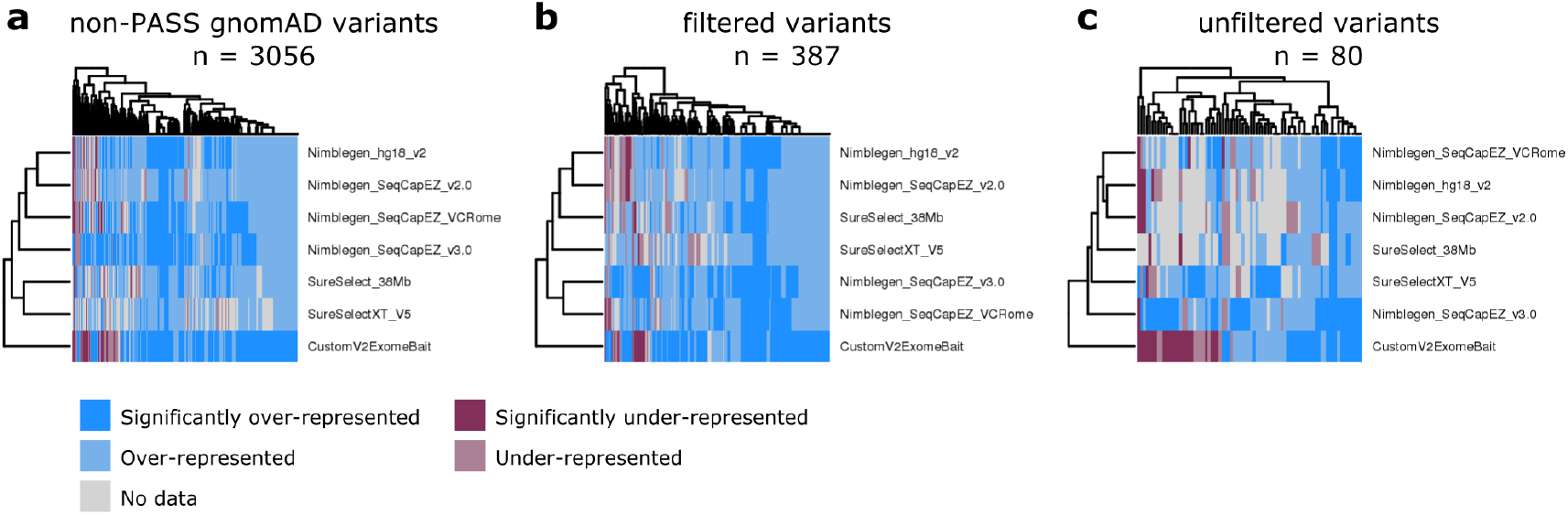
Significantly kit biased variants. Kit bias logistic regression results shown across all kits for variants that are significantly over-represented in at least one kit, as determined by linear regression. Significance threshold: Bonferroni corrected p < 0.05. **a,** Variants that failed either gnomAD exome or genome filters. **b,** Variants that failed the initial round of binomial-based filtering of mapping and sequencing artefacts. **c,** Variants that passed all previous filters.

**Supplementary Figure 4:**
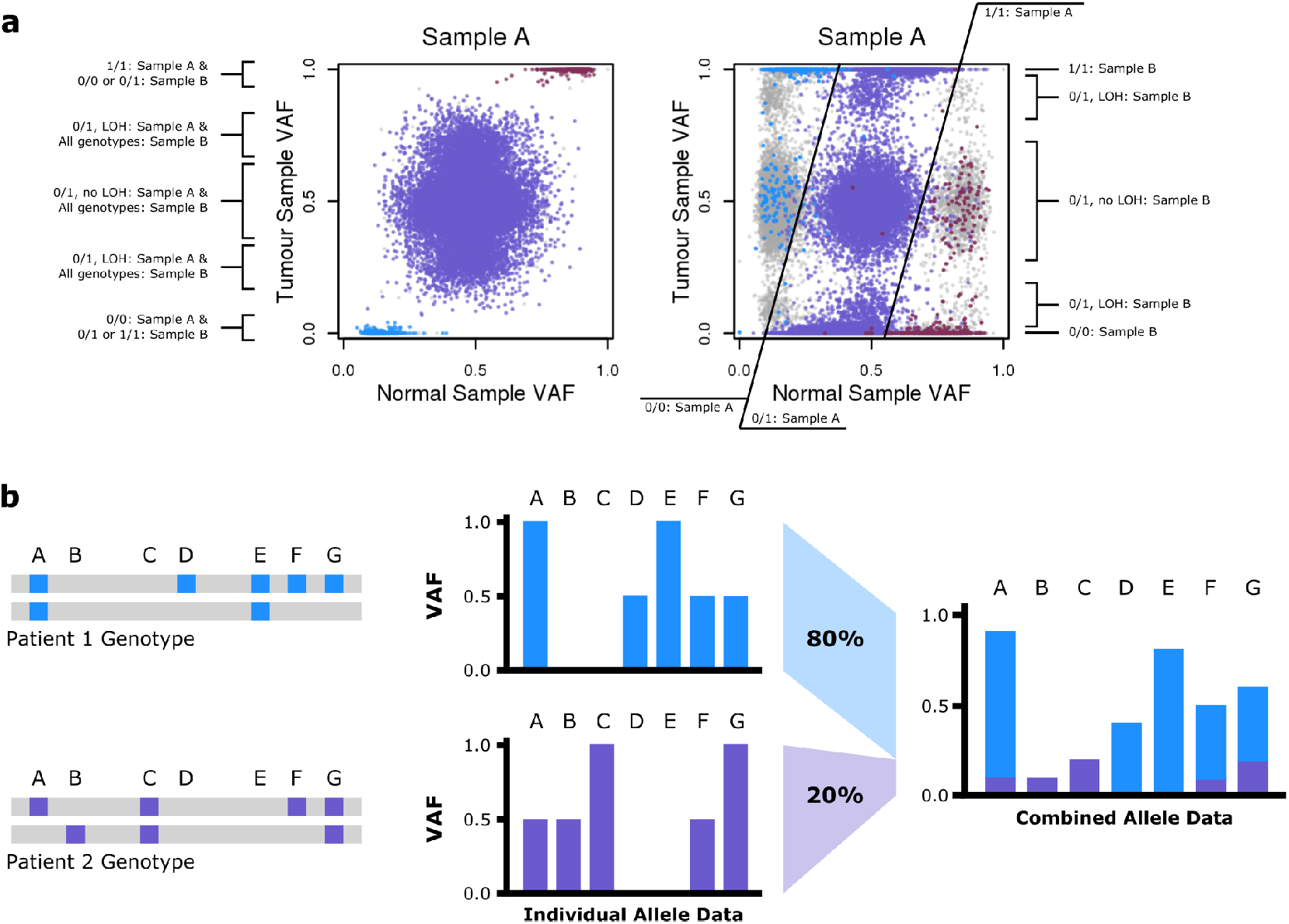
Cross contaminated patient samples in TCGA. **A**: VAF distributions of germline heterozygous variants in matched tumour:normal sample pairs from two cross-contaminated samples. Grey variants are unique to a single sample; coloured variants are shared between both samples, with colours matched between the two plots. Labels indicate the approximate limits of the clusters corresponding to the different genotype combinations. See Supplementary Table 2 for associated sequencing metadata. **B**: Schematic illustrating mixing of genotypes from two individuals at different proportions, resulting in the VAF distribution observed in **A**.

**Supplementary Figure 5:**
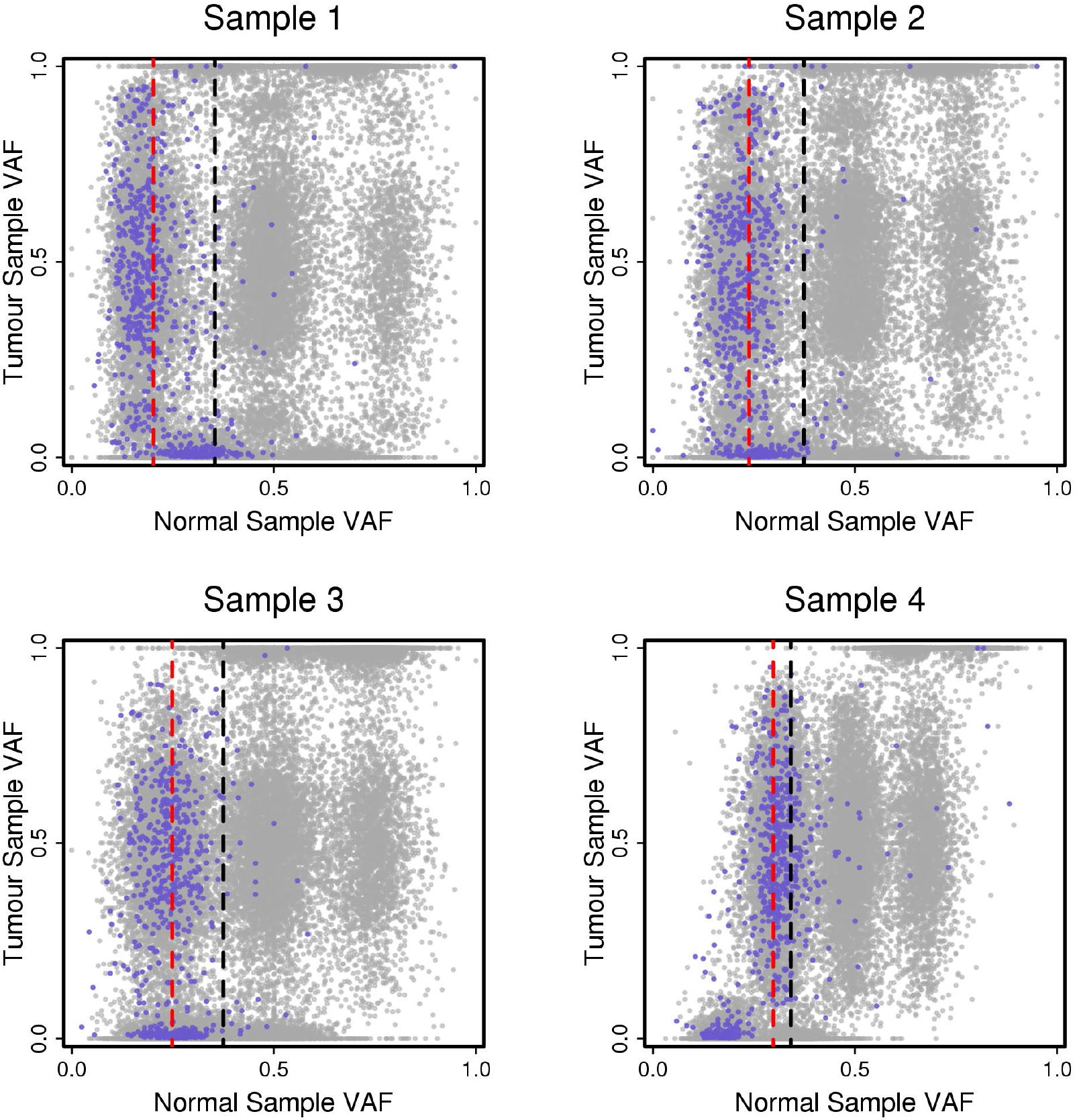
Samples processed in parallel with 50% normal sample contamination. VAF distributions of germline heterozygous variants in matched tumour:normal sample pairs from four heavily contaminated patients, sequenced in parallel. Rare variants are shown in purple, common variants in grey. Vertical dotted lines show the median VAF in normal samples for rare (red) and common (black) alleles. See Supplementary Table 3 for associated sequencing metadata.

**Supplementary Figure 6:**
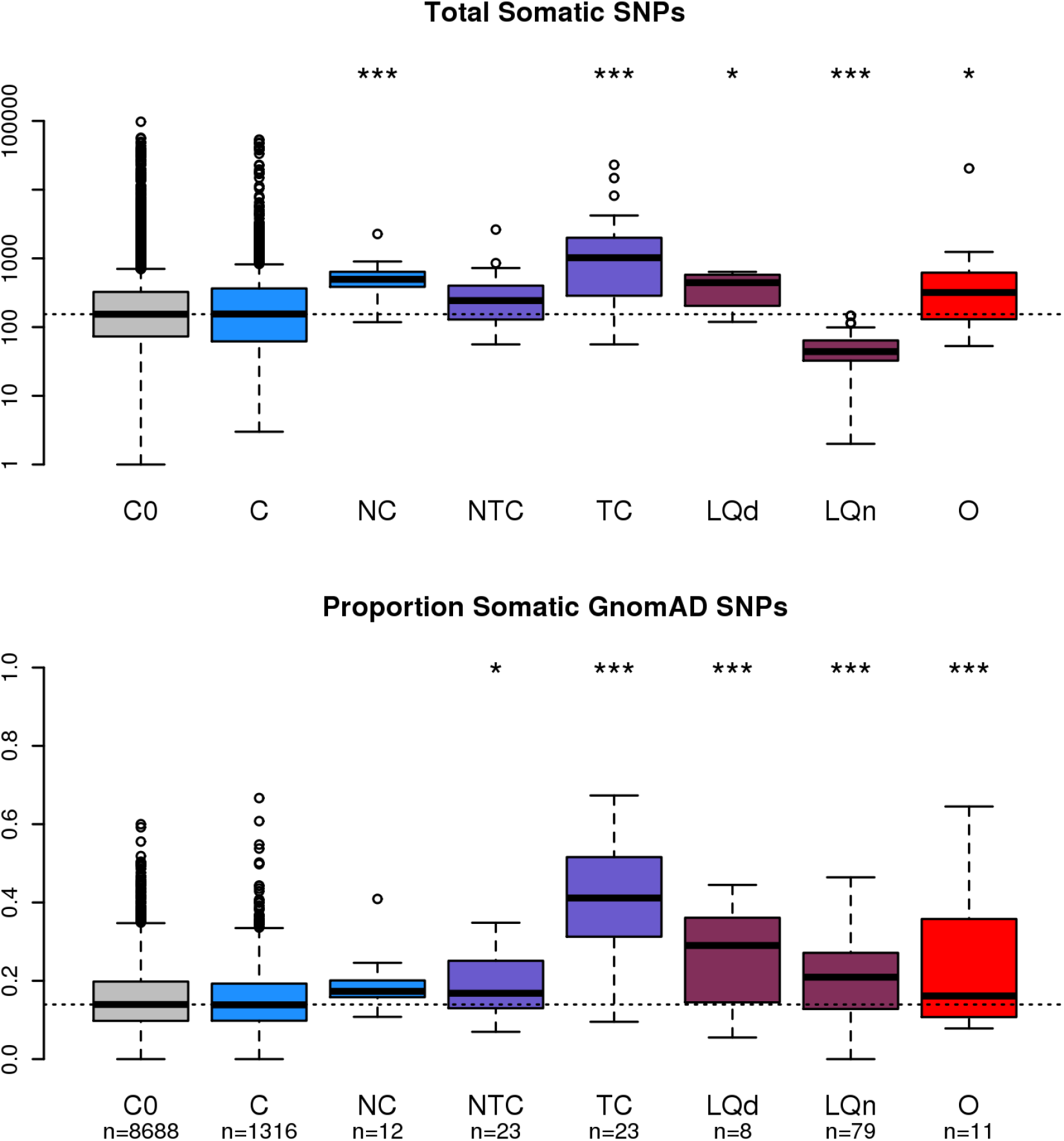
Influence of contamination and sequencing quality on somatic mutation data. Boxplots show distribution of (**A**) total somatic SNPs and (**B**) proportion of somatic SNPs found in the gnomAD database for tumour:normal sample pairs within each filtering sub-classification. Details of each sub-classification are in Table 2. Values at the bottom of the plot indicate total number of sample pairs in each group. Significance calculated by (**A**) Mann-Whitney U test compared to ‘C0’; (**B**) linear regression, response variable = proportion of somatic SNPs found in the gnomAD database, predictor variables = total somatic SNPs and filtering sub-classification. * = p-value < 0.05, ** = p-value < 0.01, *** = p-value < 0.001.

**Supplementary Figure 7:**
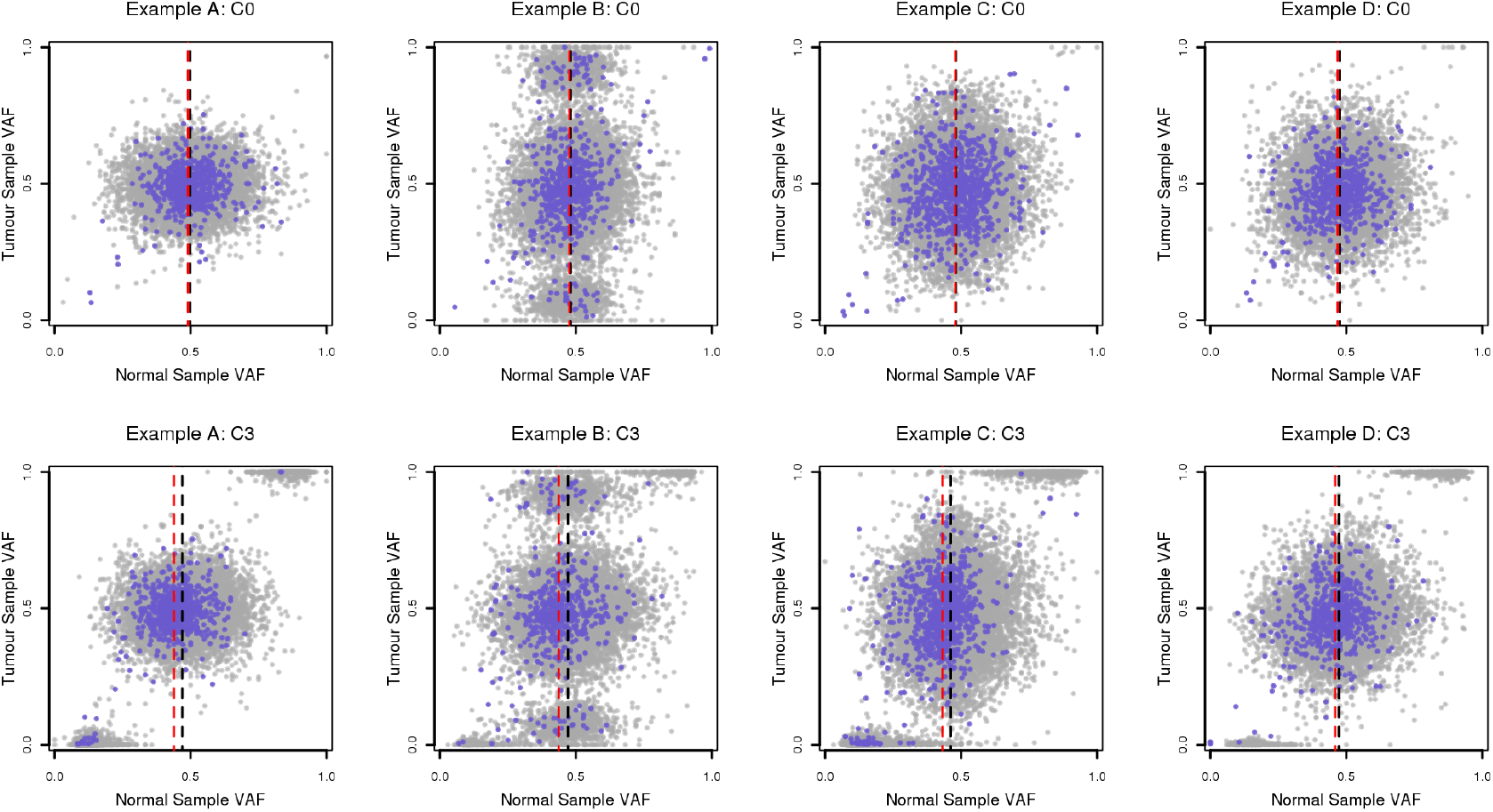
Variable degrees of contamination in pairs of samples from the same patient. VAF distributions of germline heterozygous variants in matched tumour:normal sample pairs from four patients with multiple normal samples. Within each pair, the normal samples are matched against the same tumour sample. Uncontaminated samples (C0) are on the top row, contaminated samples (C3) on the bottom row. Rare variants are shown in purple, common variants in grey. Vertical dotted lines show the median VAF in normal samples for rare (red) and common (black) alleles.

**Supplementary Figure 8:**
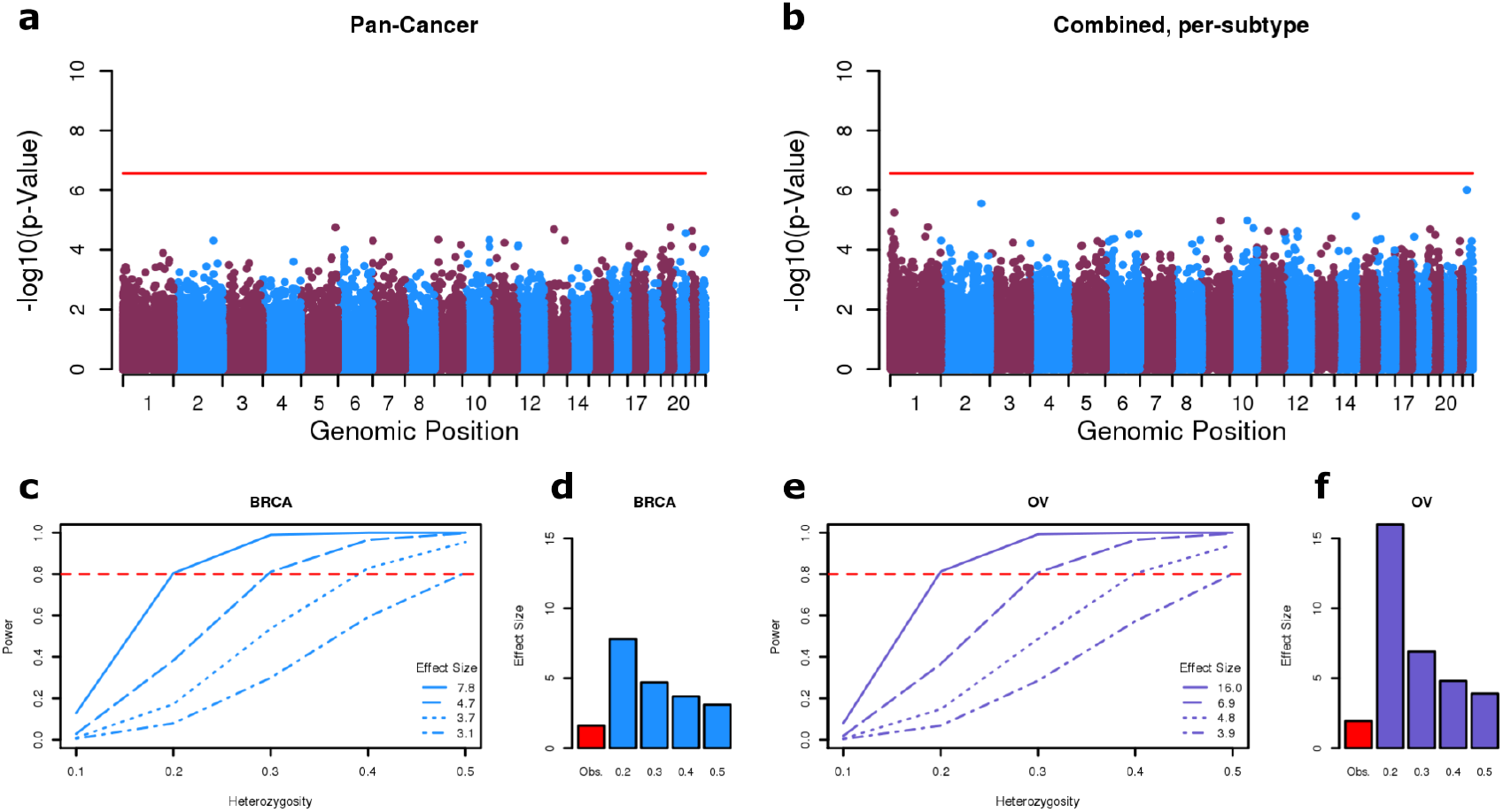
Whole-exome analysis of LOH bias. **a,** Results of the pan-cancer LOH bias analysis, post-filtering. Y-axis shows the −log10(p-value) from a Fisher’s exact test for LOH bias. Red line indicates Bonferroni corrected threshold for significance (p < 2.7e-07). **b,** Results of the per cancer subtype LOH bias analysis, post-filtering. Plot shows the most significant result across all subtypes for each variant. **c,** Results of power simulations performed for the largest cohort (breast invasive carcinoma [BRCA]) Simulations were performed using the total number of patients (831) and median frequency of LOH (0.31), and a range of heterozygosity and effect size. **d,** Barplot comparing the maximum observed GWAS effect size for BRCA (1.6), with the minimum effect size required to achieve 80% power for the indicated heterozygosity. **ef,** As in **cd**, but performed for the cohort with the highest rate of LOH (ovarian serous cystadenocarcinoma [OV]). Total number of patients = 398, median frequency of LOH = 0.45, maximum observed GWAS effect size = 1.93 (EBI GWAS Catalog).

**Supplementary Figure 9:**
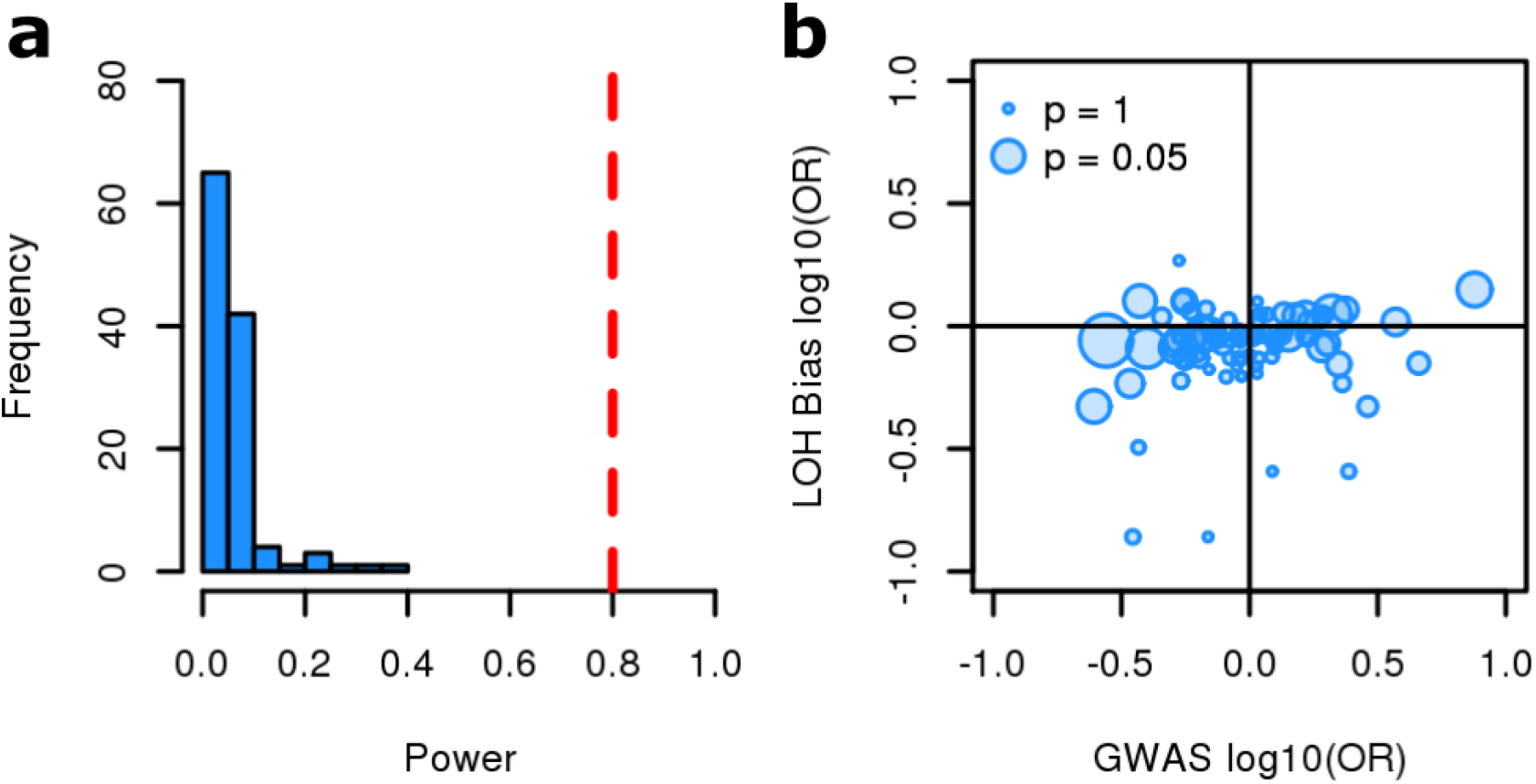
Preferential allelic retention of cancer subtype specific GWAS variants. **A**: Power to detect preferential allelic retention of cancer subtype specific GWAS variants in TCGA. Power represents proportion of simulations where p-value < 0.05. Simulations were performed using the reported GWAS OR (EBI GWAS Catalog), the total number of patients and median rate of LOH in the matched cancer subtype. **B**: Comparison of OR reported by the EBI GWAS Catalog, and the OR from the LOH bias analysis. Point sizes represent the significance of the LOH bias analysis. Full results in Supplementary Table 4.

## Supplementary Tables

**Supplementary Table 1:** Contamination classifications of all matched tumour:normal sample pairs in TCGA. <Additional File: perSamplePair.contaminationPrediction.csv>

|  | Patient UUID | BAM UUID | Sample Type | Sequencing Date<sup>1</sup> | Shared / Total Variants (%)<sup>2</sup> |
| --- | --- | --- | --- | --- | --- |
| A | 54d21956-25e4-42df-adbe-6907721fc4b5 | 085dc201-7c19-4652-a199-5daca6b1c552 | Normal | 2015-03-20T00 | 26256 / 27577 (95.2%) |
|  |  | d9bd04f9-42b6-4dbd-a770-0fa5da681290 | Tumour | 2015-01-16T00 | 22073 / 23807 (92.7%) |
| B | f8970455-bfb2-4b1d-ab71-3c5d619898ad | 089c6901-5fe6-48b0-97ab-39f00609255c | Normal | 2015-01-17T00<br>2015-01-18T00 | 26256 / 40177 (65.3%) |
|  |  | 546ba1f1-7e16-4701-875a-8e9dd426fb76 | Tumour | 2015-01-16T00<br>2015-01-18T00 | 22073 / 34019 (64.9%) |

**Supplementary Table 2:** Cross contaminated patient samples in TCGA. Associated sequencing metadata for two cross-contaminated samples from TCGA. Patient VAF distributions shown in Supplementary Figure 4a. UUID: Universally unique identifier. ^1^ GDC API endpoint: *analysis.metadata.read_groups.sequencing_date* ^2^ Italics: before filtering; plain text: after filtering. Variants = germline heterozygous variants only.

|  | Patient UUID | BAM UUID | Sample Type | Sequencing Date<sup>1</sup> |
| --- | --- | --- | --- | --- |
| 1 | 60778da8-d99e-4e51-96a2-3e900b3978d9 | 5ab3a14f-0376-49bf-93bb-3db8f5152591 | Normal | 2010-08-24T19<br>2010-08-30T19<br>2010-09-14T19 |
|  |  | 65600d4b-5164-4ae2-a520-9acdc3209a26 | Tumour | 2010-08-24T19<br>2010-08-30T19<br>2010-09-14T19 |
| 2 | 866de12b-e4df-4f19-b62e-e3fd85d4ec08 | 95167e4f-db67-40a5-86c9-9a16820538cf | Normal | 2010-08-24T19<br>2010-08-30T19<br>2010-09-14T19 |
|  |  | 35cae7a5-f598-4415-82d1-706b0ae44cec | Tumour | 2010-08-24T19<br>2010-08-30T19<br>2010-09-14T19 |
| 3 | 8e00e7e7-ffaf-44f0-91a7-172671f18e08 | f9a83975-e7b9-473a-9f8c-e398348e54fc | Normal | 2010-08-24T19<br>2010-08-30T19<br>2010-09-14T19 |
|  |  | 79a4be20-fb85-450d-9b44-f33bfdb298f8 | Tumour | 2010-08-29T19<br>2010-08-30T19<br>2010-09-14T19 |
| 4 | d846228f-67af-4b3b-9796-dbc263c2054c | cad67cb1-39df-4576-9d82-cec6605a180b | Normal | 2010-08-24T19<br>2010-08-30T19<br>2010-09-14T19<br>2010-10-23T19 |
|  |  | 454157b3-78b8-4da4-bf74-b45addabcb85 | Tumour | 2010-08-24T19<br>2010-08-30T19<br>2010-09-14T19 |

**<u>Supplementary Table 3: Samples processed in parallel with 50% normal sample contamination</u>**

Associated sequencing metadata for four highly contaminated samples from TCGA. Patient VAF distributions shown in Supplementary Figure 5.

UUID: Universally unique identifier.

^1^ GDC API endpoint: *analysis.metadata.read_groups.sequencing_date*

**<u>Supplementary Table 4: Results of cancer subtype specific GWAS variant LOH bias analysis</u>**

<Additional File: gwas.perCohort.results.csv>

**<u>Supplementary Table 5: Results of the COSMIC gene burden LOH bias analysis</u>**

<Additional File: cosmicBurden.byGene.csv>

**Supplementary Table 6:** Software and packages.

| Software / Package Name | Version |
| --- | --- |
| R | 3.3.2 |
| Perl | 5.24.0 |
| mySQL | 5.5.60-MariaDB |
| matlab | R2015b |
| gsutil | 4.28 |
| SAMtools<sup>40</sup> | 1.4.1 |
| BCFtools<sup>41</sup> | 1.3.1 |
| VCFtools<sup>42</sup> | 0.1.13 |
| BEDtools<sup>37</sup> | 2.25.0 |
| VEP<sup>22</sup> | 88 |
| gdc-client | 1.3.0 |
| Strelka<sup>13</sup> | 2.8.3 |
| CloneCNA<sup>15</sup> | 2.0 |
| GATK<sup>14</sup> | 3.8.0 |
| picard | 2.9.4 |
| optparse (R package) | 1.6.4 |
| MASS (R package) | 7.3.45 |
| scales (R package) | 0.4.1 |
| dplyr (R package) | 0.5.0 |
| RColorBrewer (R package) | 1.1.2 |

**Supplementary Table 7:** Exome target capture kit genomic region files.

| Kit | Source | File Name |
| --- | --- | --- |
| Nimblegen_2.1M | Nimblegen | 2.1M_Human_Exome.capture.hg19.bed |
| Nimblegen_hg18_v2 | Nimblegen | hg18_nimblegen_exome_version_2.bed |
| Nimblegen_SeqCapEZ_v2.0 | Nimblegen | SeqCap_EZ_Exome_v2.bed |
| Nimblegen_SeqCapEZ_v3.0 | Nimblegen | SeqCap_EZ_Exome_v3_capture.bed |
| Nimblegen_SeqCapEZ_VCRome | Nimblegen | VCRome_2_1_hg19_primary_targets.bed |
| SureSelect_38Mb | Agilent | S0293689_Covered.bed |
| SureSelectXT_V5 | Agilent | S04380110_Covered.bed |
| CustomV2ExomeBait | Agilent | whole_exome_agilent_1.1_refseq_plus_3_boost<br>ers.targetIntervals.bed |

**Supplementary Table 8:** Collapsed ClinVar Categories.

|  |  |
| --- | --- |
| **Benign** |  |
| Benign | Benign/Likely_benign,_risk_factor |
| Likely_benign | Benign/Likely_benign,_protective |
| Benign/Likely_benign | Benign/Likely_benign,_association |
| Benign/Likely_benign,_other |  |
| **Pathogenic** |  |
| Pathogenic | Pathogenic,_risk_factor |
| Pathogenic/Likely_pathogenic | Pathogenic/Likely_pathogenic,_risk_factor |
| Likely_pathogenic | Pathogenic,_Affects |
| Likely_pathogenic,_risk_factor |  |
| **Unknown** |  |
| Uncertain_significance | Conflicting_interpretations_of_pathogenicity,_risk_factor |
| not_provided | Conflicting_interpretations_of_pathogenicity |
| drug_response | drug_response,_protective,_risk_factor |
| risk_factor | . |
| **Missing** |  |
| *Not in ClinVar database* |  |

## Notes

https://git.ecdf.ed.ac.uk/taylor-lab/GenomeArtiFinder

https://doi.org/10.7488/ds/2860

### Competing Interest Statement

The authors have declared no competing interest.

